# High-Complexity Barcoded Rabies Virus for Scalable Circuit Mapping Using Single-Cell and Single-Nucleus Sequencing

**DOI:** 10.1101/2024.10.01.616167

**Authors:** David Shin, Madeleine E. Urbanek, H. Hanh Larson, Anthony J. Moussa, Kevin Y. Lee, Donovan L. Baker, Elio Standen-Bloom, Sangeetha Ramachandran, Derek Bogdanoff, Cathryn R. Cadwell, Tomasz J. Nowakowski

**Author notes:** These authors contributed equally (David Shin, Madeleine E. Urbanek). Senior author (Cathryn R. Cadwell, Tomasz J. Nowakowski).

## Abstract

Single cell genomics has revolutionized our understanding of neuronal cell types. However, scalable technologies for probing single-cell connectivity are lacking, and we are just beginning to understand how molecularly defined cell types are organized into functional circuits. Here, we describe a protocol to generate high-complexity barcoded rabies virus (RV) for scalable circuit mapping from tens of thousands of individual starter cells in parallel. In addition, we introduce a strategy for targeting RV-encoded barcode transcripts to the nucleus so that they can be read out using single-nucleus RNA sequencing (snRNA-seq). We apply this tool in organotypic slice cultures of the developing human cerebral cortex, which reveals the emergence of cell type– specific circuit motifs in midgestation. By leveraging the power and throughput of single cell genomics for mapping synaptic connectivity, we chart a path forward for scalable circuit mapping of molecularly-defined cell types in healthy and disease states.

## INTRODUCTION

Understanding neuronal connectivity at the cell type level is crucial for unraveling the complexities of brain function and dysfunction. Monosynaptic tracing using glycoprotein (G)–deleted rabies virus (RVdG) has been a powerful method for mapping direct synaptic connections between populations of cells^1,2^. This method exploits the natural ability of RVdG to spread trans-synaptically in the retrograde direction, allowing researchers to trace connections from a genetically targeted starter cell population to its presynaptic partners. Despite its utility, traditional monosynaptic rabies tracing methods have limitations. For example, these techniques typically offer only coarse cell-type specificity due to the limitations of transgenic mouse lines and viral tools used for targeting specific cell populations. With the advent of single-cell RNA sequencing, the neuronal landscape has been shown to be much more complex, revealing a multitude of fine-grained cell types that cannot be distinguished with existing tracing methods.

Several recent studies have incorporated a nucleotide ‘barcode’ into the RVdG backbone, enabling simultaneous readout of the connectivity and cell identity of hundreds of cells in a single experiment using single cell or spatial transcriptomics assays^3–5^. However, this barcoded RVdG technology still faces significant challenges. First, in contrast to other viruses such as lentivirus or adeno-associated virus, it has been challenging to achieve high-complexity libraries of barcoded RVdG, and the limited diversity of barcoded RVdG libraries represents a major roadblock to scaling this technology to study larger networks. There are multiple reasons for this, including the low efficiency of active viral RNA particle recovery from a DNA template and variable clonal expansion of the few barcodes that are recovered, among others^6–8^. Second, given that RVdG replicates and undergoes transcription in the cytoplasm^9,10^, it has not been possible to read out RVdG barcodes from isolated nuclei, necessitating whole cell dissociations which compromise neuron viability and integrity, further limiting the utility of this tool.

To overcome these challenges, we have developed an optimized protocol for RVdG packaging to improve the complexity of barcoded RVdG libraries by several orders of magnitude compared to currently published tools. This advancement significantly enhances the scalability of trans-synaptic tracing experiments, allowing us to potentially map the inputs to tens of thousands of single cells in parallel. Additionally, we have incorporated a method for localizing RVdG barcodes to nuclei, allowing for high-fidelity connectivity mapping without the need for whole cell dissociations. To demonstrate the power of this tool, we characterize the emergence of cell type–specific connectivity patterns by mapping the inputs to thousands of single starter cells in organotypic slices of the developing human cerebral cortex.

## RESULTS

### Generation of high-complexity barcoded RVdG libraries enables mapping the inputs to thousands of single cells in parallel

To generate a high-complexity barcoded RVdG plasmid library, we have leveraged a recently developed combinatorial barcode system optimized for scRNA-seq readout^11^. This barcode ‘white list’ system generates barcode complexity exponentially using combinatorial indexing, while maintaining a high (≥6) minimum Hamming distance that is robust to sequencing errors (Figures S1A and S1B; Table S1). We cloned these barcodes into the 3’ UTR of the fourth open reading frame of the RVdG genome plasmid encoding dTomato (Figures 1A, 1B, S1C and S1D; Table S2).

**Figure 1.**
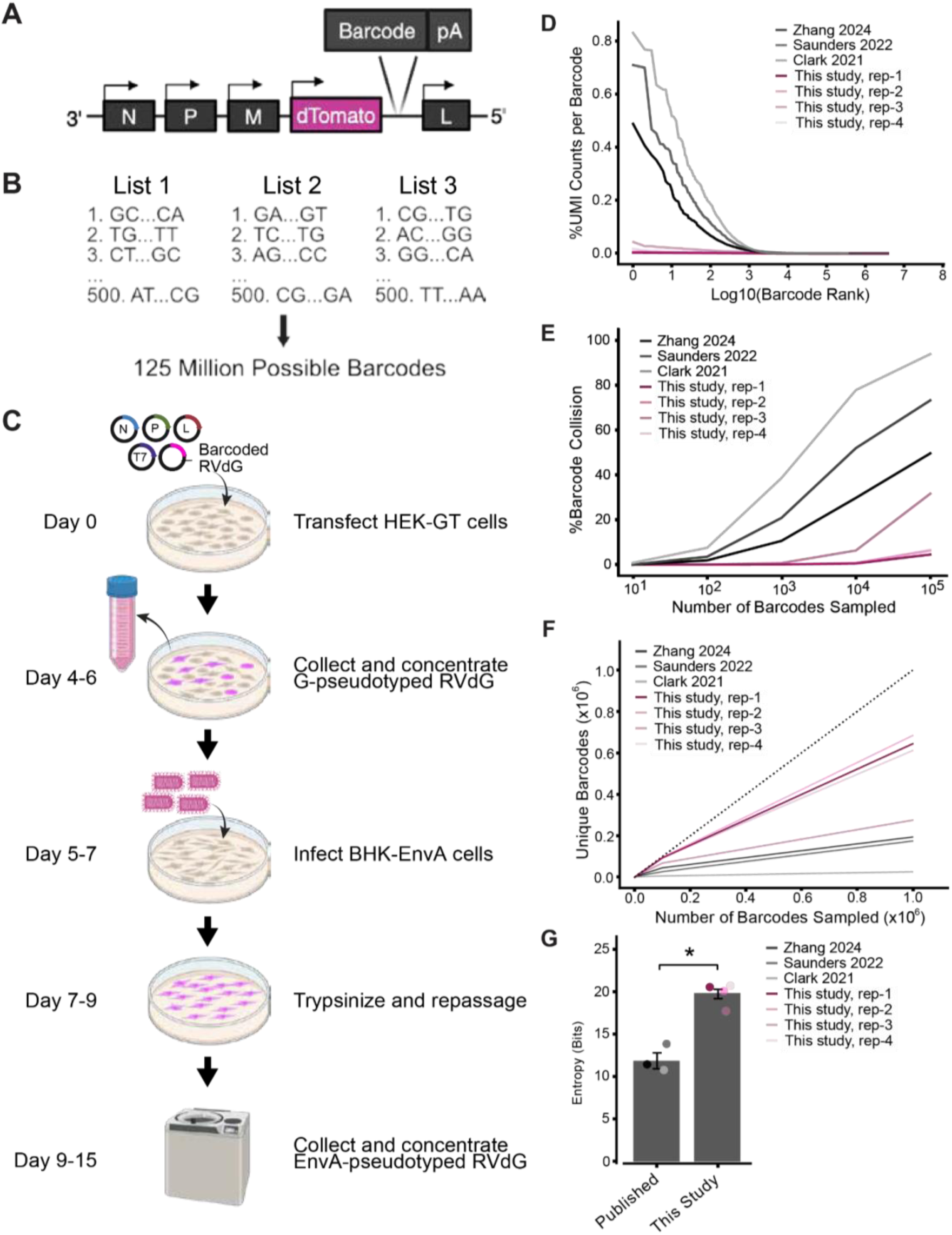
Generation of high-complexity barcoded RVdG libraries. **A.** Location of barcode cloning site in the rabies genome. **B**. Design of whitelist combinatorial barcode system. **C.** Schematic of barcoded RV production protocol. **D.** Distribution of UMI counts per barcode normalized as a percent of total barcode UMI counts. Barcoded RVdG libraries from this study are compared to previously published libraries^3–5^. **E**. Percentage of simulated barcode collision events, or two cells receiving the same barcode by chance, as a function of the number of infections simulated. Error bars are 95% confidence intervals for 50 iterations. **F.** Number of unique barcodes identified as a function of sequencing depth. Error bars are standard deviation for 50 iterations. **G.** Shannon entropy (in bits) of each viral library. P-value computed using the Student’s t-test (*p<0.05; published: n = 3 independent libraries, this study: n = 4 independent libraries [replicates 1 and 2 = SAD B19, replicates 3 and 4 = CVS-N2c]).

We optimized a protocol for packaging RVdG to improve barcode complexity and promote a more uniform distribution of viral barcodes (Figure 1C–1G and Figure 2). Using this optimized RVdG packaging protocol, we were able to generate high-complexity barcoded RVdG libraries which contained millions of unique barcodes with a relatively even barcode distribution compared to prior published libraries^3–5^ (GEO: GSE277536). Specifically, no single barcode in our packaged RVdG libraries accounts for more than 0.046% of the total number of barcodes detected (Figure 1D). In order to estimate the number of unique starter infections that can be traced using different barcoded RVdG libraries, we used a simulation to compute the rate of barcode collision, or at least two starter cells receiving the same barcode, as a function of the number of cells infected. This simulation revealed that our optimized RVdG libraries enable unique labeling of 10,000 to 100,000 single cells with a barcode collision rate less than 10%, which represents two to three orders of magnitude improvement compared to previously published libraries (Figure 1E). This increased barcode complexity of our libraries persisted across a broad range of simulated sequencing depths (FIgure 1F). We also quantified the complexity of our barcoded RVdG libraries using an estimate of Shannon entropy^12,13^ and found that our optimized libraries correspond to an approximately 8-bit improvement in information content compared to previously published libraries (Figure 1G). A difference in a single bit represents a doubling of library complexity through increasing either the number of barcodes, uniformity of the barcode distribution, or both (STAR Methods). In summary, our optimized RVdG packaging protocol leads to a robust improvement in barcoded RVdG library complexity compared to prior studies.

**Figure 2.**
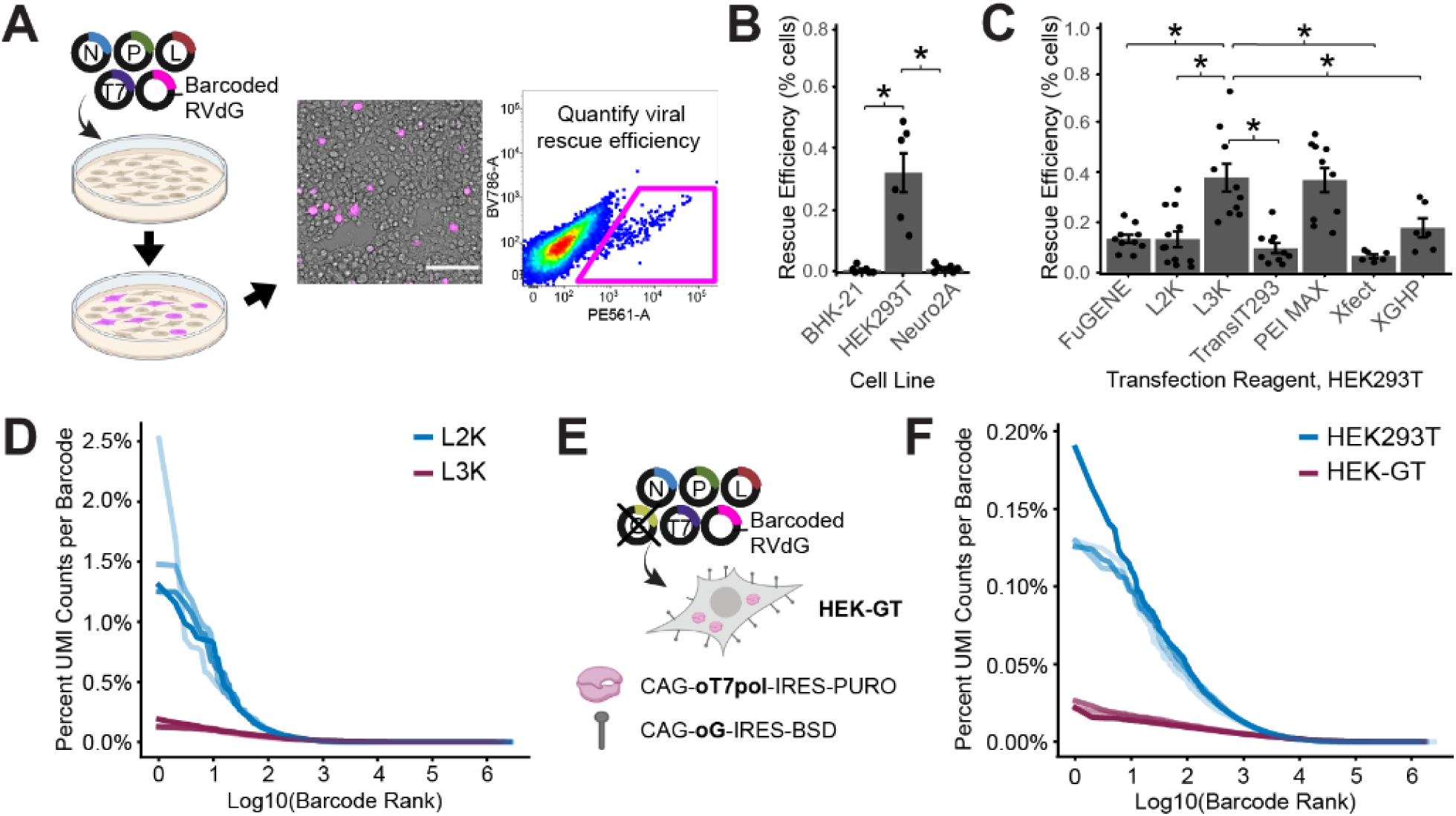
Viral packaging cell lines and transfection reagents are critical to generate high-complexity barcoded RVdG. **A.** To assess packaging efficiency, cells are transfected with RVdG packaging plasmids in the absence of rabies G, and the percentage of cells producing rabies virus three days after transfection is measured by flow cytometry. **B.** Percentage of cells successfully producing RVdG three days after transfection using Lipofectamine 3000 (Lipo3000), comparing three commonly used packaging cell lines: BHK-21 (n=6 technical replicates), HEK293T (n=6 technical replicates), and Neuro2A (n=6 technical replicates). **C.** Percentage of HEK293T cells successfully packaging RVdG three days after transfection, comparing various commonly used transfection reagents: FuGENE HD (n = 10 technical replicates), Lipofectamine 2000 (n = 12 technical replicates), Lipofectamine 3000 (n = 10 technical replicates), TransIT-293 (n = 10 technical replicates), PEI (n = 10 technical replicates), Xfect (n = 6 technical replicates), and XtremeGene HP (n = 6 technical replicates). **D.** Distribution of UMI counts per barcode normalized as a percent of total UMI counts, comparing libraries generated from HEK293T cells using Lipo3000 (n=4 technical replicates) and Lipofectamine 2000 (Lipo2000, n=4 technical replicates). Transfection for virus packaging included rabies G. **E.** Schematic of virus packaging using HEK-GT cell line, which was engineered to stably express human codon–optimized T7 polymerase (oT7pol) and the optimized rabies glycoprotein (oG), requiring fewer plasmids to be co-transfected during packaging. **F.** Distribution of UMI counts per barcode normalized as a percent of total UMI counts, comparing libraries generated in HEK293T cells (n=4 technical replicates) compared to HEK-GT cells (n=4 technical replicates) using Lipofectamine 3000. In **(B)** and **(C)**, p-values are computed using Wilcoxon Rank Sum with Bonferroni-Holm correction for multiple comparisons, *p<0.05.

### The HEK-GT cell line and and Lipofectamine 3000 transfection reagent are optimal for generating high-complexity barcoded RVdG libraries

Current state-of-the-art protocols for packaging RVdG involve a three-step process^4,6,14,15^: First, a packaging cell line (e.g. HEK293T, Neuro2A, or B7GG cells) undergoes transfection with packaging plasmids, which include an RVdG genome plasmid, four plasmids encoding RV proteins (N, P, L, and G) necessary for RVdG packaging and propagation, and a plasmid encoding T7 polymerase to drive the expression of the RVdG genome. The initial recovery of replication-competent viral particles from plasmid DNA at this step is extremely inefficient, with some reports suggesting that less than 0.01% of transfected cells successfully begin to produce RVdG^4,16^. As a result, in the second step RVdG is given time to propagate throughout the culture until a majority of cells are producing the virus (Figure S1E). Third, G-pseudotyped viral supernatant is harvested and used to transduce BHK21 cells that stably express EnvA in order to produce EnvA-pseudotyped RVdG (Figure S1F), which is subsequently collected and concentrated by ultracentrifugation and used for downstream experiments.

We reasoned that improving the efficiency of recovering viral particles from plasmid DNA might improve the diversity and distribution of our barcoded RVdG viral libraries (Figure 2A). We evaluated the efficiency of commonly used cell lines for packaging RVdG: HEK293T, BHK-21, and Neuro2A (Figures 2B, S1G–S1J). We transfected each cell line with packaging plasmids in the absence of rabies G to disable cell-to-cell viral spread, and quantified the percentage of cells that rescued RVdG using flow cytometry. HEK293T cells were more efficient at packaging RVdG, rescuing RVdG at rates approximately 48-fold higher than BHK-21 cells and ∼29-fold higher than Neuro2A cells (HEK293T: 0.32±0.13%, BHK21: 0.0067±0.0027%, Neuro2A: 0.011±0.0045%, mean±SEM; HEK293T vs. BHK21: p=0.013, HEK293T vs. Neuro2A: p= 0.013, BHK21 vs. Neuro2A: p = 0.53, Wilcoxon rank sum test with Bonferroni-Holm correction for family-wise error rate; Figures 2B, S1G– S1J).

We also tested different transfection reagents to identify those that might further improve viral rescue efficiency in HEK293T cells (Figures 2C, S1I, and S1J). Lipofectamine 3000 (Lipo3000) and PEI MAX showed the highest efficiencies of viral rescue (Lipo3000: 0.38±0.05%; PEI MAX: 0.37±0.05%). These reagents yielded a 2.9-fold improvement in viral rescue on average compared to Lipofectamine 2000 (Lipo2000) and a 5.8-fold improvement compared to Xfect, two transfection reagents previously used for barcoded RVdG packaging^3–5^ (Figures 2C, S1I, and S1J).

We next prepared barcoded viral libraries to directly compare the diversity achieved using Lipo3000 and Lipo2000 (STAR Methods) in HEK293T cells. RVdG prepared with Lipo3000 exhibited a more even barcode distribution compared to Lipo2000, with the most common barcode in Lipo3000 libraries accounting for on average 0.18% of total barcoded genomes detected, compared to 2.84% in Lipo2000 libraries (Figure 2D). The rate of barcode collision was lower, and the Shannon entropy was higher, in Lipo3000 compared to Lipo2000 libraries (Figures S1K and S1L; STAR Methods).

We reasoned that some barcoded RVdG particles might gain a competitive advantage during viral propagation due to variability in rabies G expression, leading to overrepresentation of a subset of barcodes. To test this, we leveraged a recently developed packaging cell line, HEK-GT, which is a HEK293T cell line that stably expresses the chimeric rabies glycoprotein oG and T7 polymerase^17,18^ (Figure 2E). Indeed, HEK-GT cells produced RVdG libraries with a more uniform barcode distribution compared to HEK293T (Figures 2F, S1K, and S1L). Another proposed solution for uneven viral propagation during packaging is to physically constrain the amplification of barcoded RV clones by using multi-well plates with a smaller surface area^19^. However, we found no difference in library complexity between barcoded RV libraries packaged in 15 cm dishes and 384-well plates (Figures S1M-S1N).

### Helper virus– and RVdG–infected cells are predominantly neuronal in human prenatal organotypic slice cultures

To demonstrate the utility of our barcoded RVdG circuit mapping strategy in a biological circuit, we prepared organotypic slice cultures from the human cerebral cortex during the second trimester of prenatal development, spanning gestational weeks (GW) 16-22. At this developmental stage, puncta positive for the postsynaptic marker PSD-95 are enriched in the marginal zone and along structures resembling dendrites in the cortical plate, while puncta positive for the presynaptic markers synaptophysin (SYP) and VGLUT1 are concentrated in the marginal zone and the upper layer of the subplate (Figures 3A, 3B, and S2A), consistent with previous studies investigating the distribution of synapses at this age^20,21^. To perform circuit mapping using barcoded RVdG, slices were first infected with a lentivirus helper on day 0 (DIV0) which encodes an optimized RV glycoprotein (oG), which is necessary for trans-synaptic spread, TVA, an avian receptor that enables selective infection by EnvA-pseudotyped viral particles, and the fluorescent reporter GFP or mGreenLantern-KASH (mGL-KASH) (Figure 3C; STAR Methods). On DIV3, slices were transduced with RVdG expressing either dTomato or oScarlet-KASH (Figure 3C and S2B–S2H). By DIV8, we observed both double-labeled starter cells and putative presynaptic non-starter cells positive for RVdG only (Figures 3D–3G and S2B–S2H). Immunohistochemistry revealed that most of the infected cells expressed postmitotic neuronal markers NeuN and NEUROD2, suggesting that the majority of RVdG+ starter and non-starter cells are glutamatergic neurons (Figures 3D–3F and S2B–S2H).

**Figure 3.**
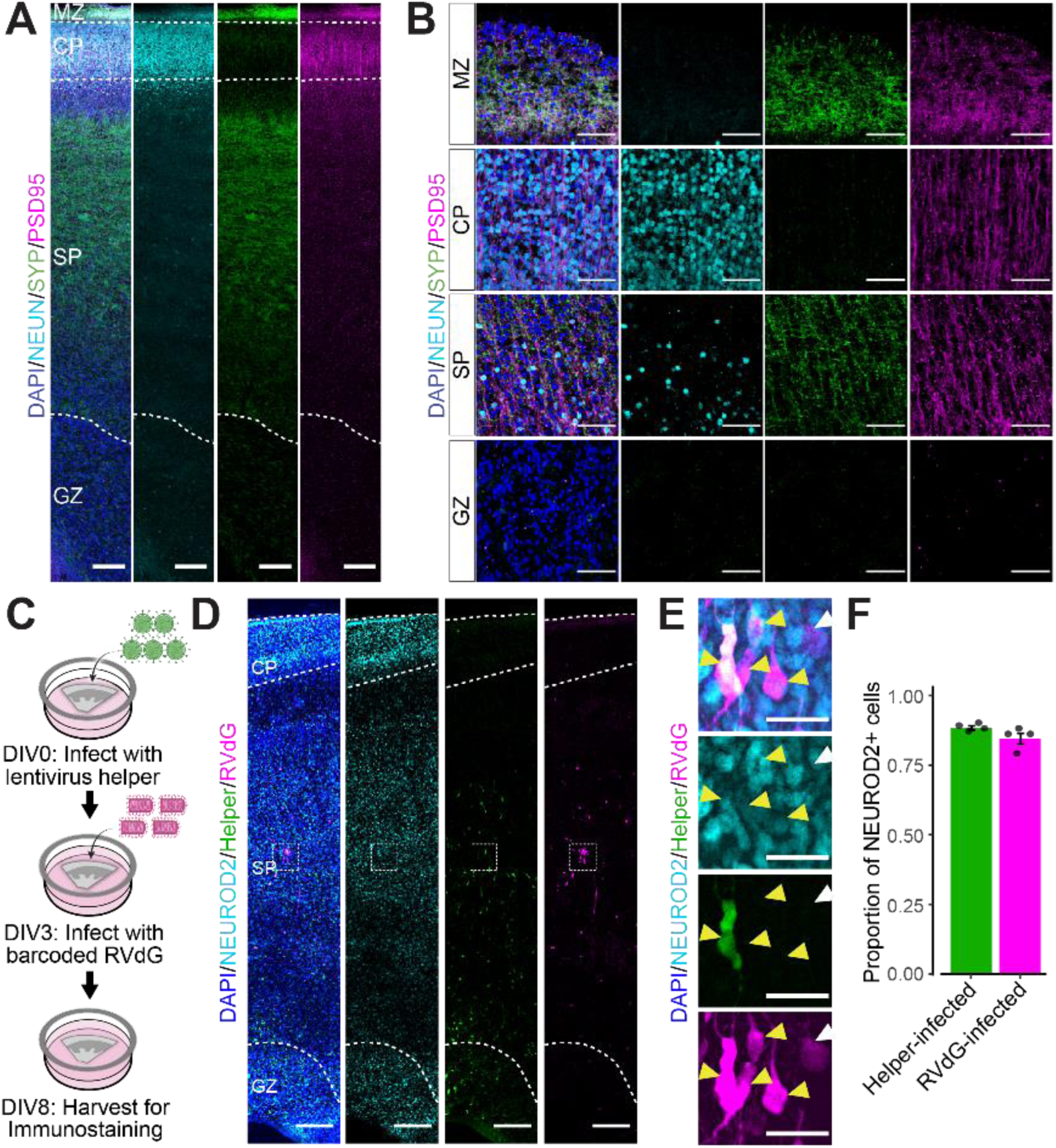
Helper virus– and RVdG–infected cells are predominantly neuronal in human prenatal organotypic slice cultures. **A-B.** Full thickness tile scan **(A)** and high magnification images **(B)** of gestational week (GW) 22 cortex stained for pan-neuronal marker NeuN, presynaptic marker synaptophysin (SYP), and postsynaptic marker PSD95. **C**. Schematic of RVdG circuit mapping experiment in human prenatal organotypic slice cultures. **D-E**. Full thickness tile scan **(D)** and high magnification image **(E)** of slice stained for GFP (cells infected with helper virus), RFP (cells infected with RVdG), and NeuroD2. **F**. Quantification of the proportion of helper virus**–** and RVdG**–**infected cells that co-express NEUROD2 (n=4 slices from 4 different donors). Scale bars: 250 μm in panels **(A,D)**, 50 μm in panel **(B),** 25 μm in panel **(E)**.

### Helper virus– and RVdG–infected cells are predominantly layer 2/3 excitatory neurons

To achieve a more granular classification of RVdG-infected cell types, we performed single-cell RNA sequencing (scRNA-seq). Tissue slices from seven individuals were dissociated into single cells and FACS-sorted based on the expression of RVdG-encoded dTomato or oScarlet-KASH (Figures 4A, S3A, and S3B). In a subset of slices, we also collected dTomato/oScarlet-negative cells to serve as uninfected controls. We captured a total of 17,343 RVdG-infected cells and 7,446 uninfected cells, after filtering based on quality control metrics (STAR Methods). We used principal component analysis to reduce the dimensionality of the data and Harmony^22^ for dataset integration. We visualized the transcriptomic data in high dimensional space using uniform manifold approximation and projection^23^ (UMAP) and used Louvain clustering^24^ to group cells into distinct clusters based on their gene expression profiles (Figure 4B). We annotated the resulting transcriptomic clusters based on expression of canonical marker genes to identify the major cell types: layer 2/3 neurons (L2/3 ExNs) that express BHLHE22^25,26^ and CUX2^27–30^, layer 4 neurons (L4 ExNs) that express BHLHE22^25,26^ and RORB^31,32^, layer 5/6 and subplate neurons positive for TLE4^30,33–36^ and CRYM^30,37–39^ (L5/6/SP ExNs), DLX5/GAD1+ inhibitory neurons (InNs)^40–42^, PDGFRA+ oligodendroglia (OL)^43,44^, EOMES+ intermediate progenitor cells (IPCs)^45–47^, HES1+ radial glia (RG)^48–50^, astrocytes enriched for AQP4^48,51–53^ and GFAP^54–56^ (AC), and MKI67+ dividing progenitors^57^ (DIV) (Figures 4C-D, S4 and Table S3). Each cell type was represented in each sample (Figure 4E). RVdG-infected cells were further divided into starter and non-starter populations based on the expression of lentiviral helper transcripts (STAR Methods). Compared to uninfected cells, RVdG-infected cells were enriched in GABAergic, L2/3, and L4 ExNs and depleted of RG and astrocytes (one-sample t-test, p=0.00088, 4.304e-09, 6.784e-05, 1.5968e-07, and 6.432e-06) (Figures 4F and 4G).

**Figure 4.**
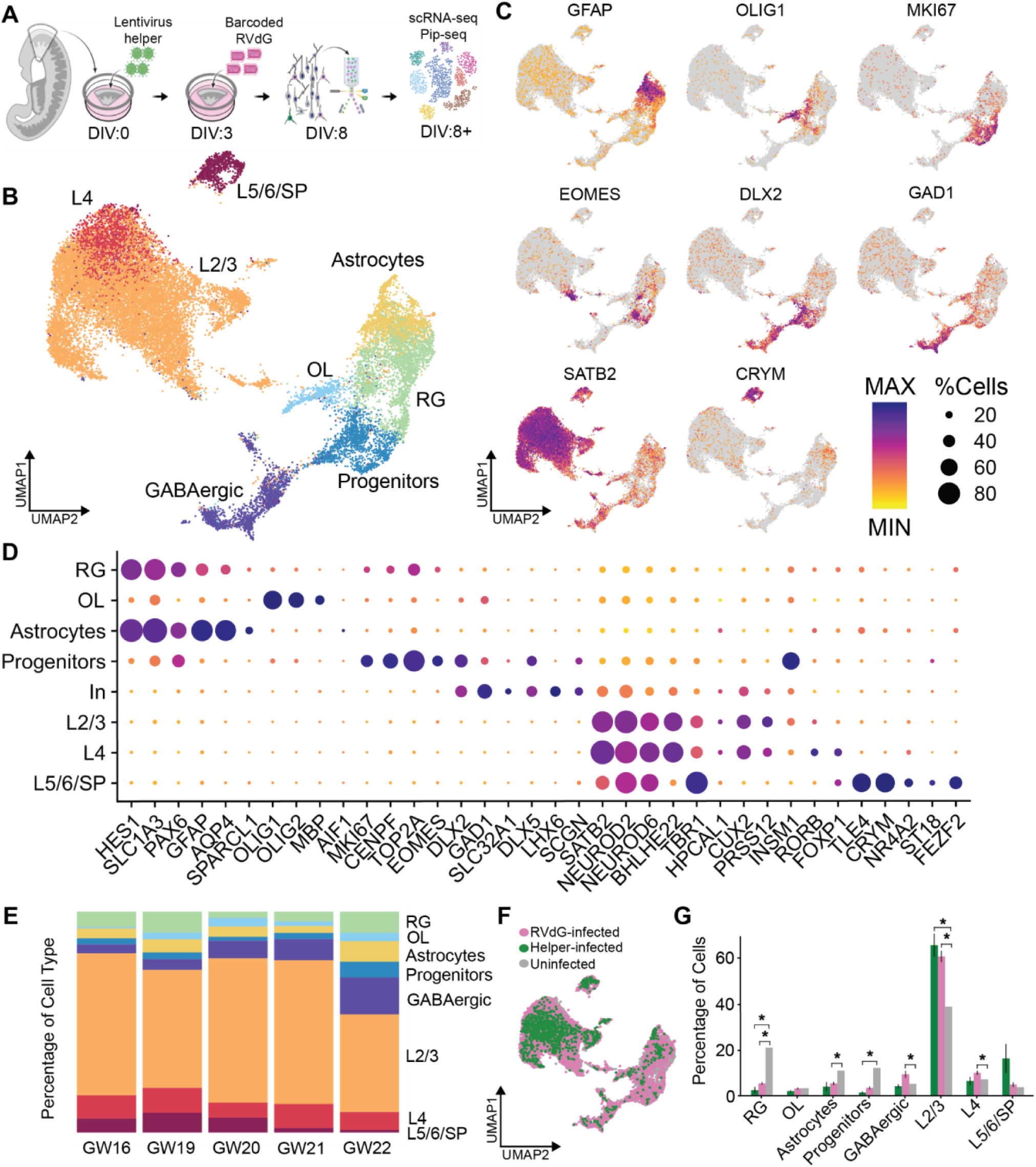
Helper virus– and RVdG–infected cells are predominantly layer 2/3 excitatory neurons. **A.** Schematic of dissociated RNA-sequencing analysis of RVdG-infected tissue. **B.** UMAP of all pooled datasets annotated with cell types. **C.** Feature plots showing expression of key cell type marker genes across clusters. **D.** Dot plot of key marker genes across cell type clusters. Color indicates relative expression and size indicates percent of expressing cells within the cluster. **E.** Stacked bar plot showing cell type distribution across sample ages. **F.** UMAP with cells colored to match either RVdG and helper virus infection status. **G.** Proportions of cell types among RVdG-infected, helper-infected or uninfected populations (bar plot indicates mean±SEM; p-values computed using one sample t-test comparing RVdG-infected vs. uninfected and helper-infected vs. uninfected and Bonferroni correction for multiple comparisons, *p<0.05; n=9 datasets for RVdG- and helper-infected populations, n=1 dataset for uninfected population).

Similarly, when compared to uninfected cells, helper-infected cells were enriched in L2/3 ExNs and depleted of RG and progenitors (one-sample t-test, p=0.00088, 0.01650912, 2.464e-05, and 3.408e-07) (Figure 4G).

### RVdG-infected neurons exhibit strain-dependent toxicity

Prior work has suggested that the CVS-N2c strain of RVdG is less toxic and spreads more efficiently across synapses compared to the more commonly used SAD-B19 strain^58^. Since our dataset includes slices infected with both the SAD-B19 strain (n=5 individuals at GW 16, 20, 20, 21, and 22) and the CVS-N2c strain (n=2 individuals at GW 19 and 21), we next set out to quantify strain-dependent toxicity profiles and cell type biases. In total, we identified 5,840 cells from SAD B19 RVdG–infected slices and 11,503 cells from CVS-N2c RVdG– infected slices after filtering based on quality control metrics (Figure 5A and STAR Methods).

**Figure 5.**
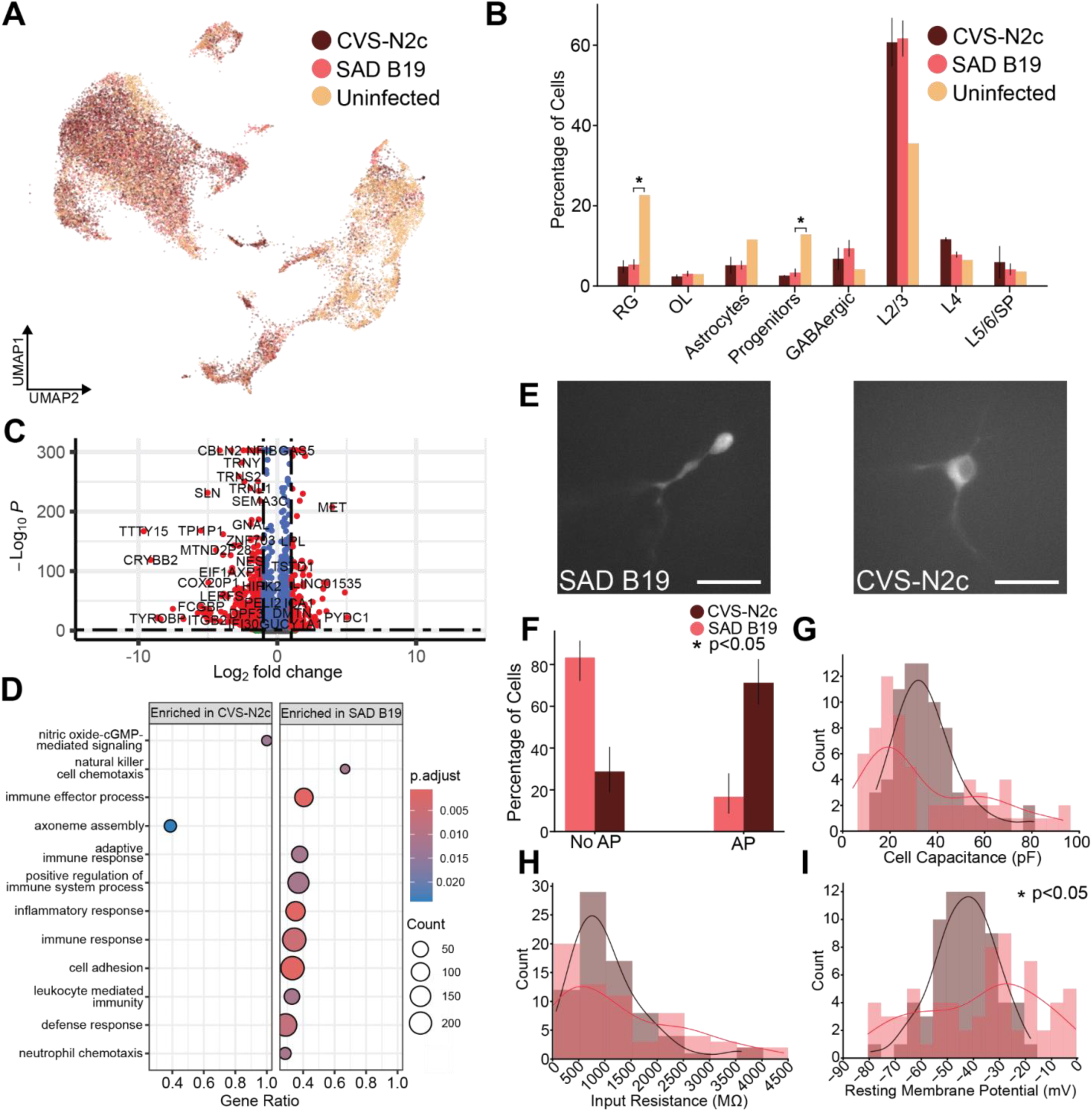
RVdG-infected neurons exhibit strain-dependent toxicity. **A.** UMAP visualization of transcriptomic cell types from SAD B19–infected, CVS-N2c–infected, and uninfected cell populations. **B.** Percentage of each major cell type among SAD B19–infected, CVS-N2c–infected and uninfected cell populations (bar plot indicates mean±SEM; *p<0.05, one sample t-tests comparing CVS-N2c versus uninfected and SAD B19 versus uninfected within each cell type, with Bonferroni correction for multiple comparisons; n=2 samples for CVS-N2c, n=5 samples for SAD B19, and n=1 sample for uninfected datasets). **C.** Volcano plot showing differentially expressed genes between CVS-N2c– and SAD B19–infected cells. Positive log_2_(fold change) values indicate enrichment in CVS-N2c–infected cells, while negative log_2_(fold change) values indicate enrichment in SAD B19–infected cells. Genes with an adjusted p-value ≤ 0.05 are shown in red. Genes from the GO term “Immune Response” are annotated. **D.** Gene set enrichment analysis reveals an enrichment of immune response pathways in SAD B19–infected compared to CVS-N2c–infected cells. **E.** Representative fluorescent images of SAD B19– and CVS-N2c–infected neurons. Scale bars, 30 μm. **F.** Percentage of infected neurons capable of firing action potentials in response to direct current injection. Error bars are 95% Clopper-Pearson confidence intervals. P-value computed using Fisher’s exact test (n=66 recordings for SAD B19, n=71 recordings for CVS-N2c). **G-I.** Intrinsic properties of recorded neurons including cell capacitance **(G)**, input resistance **(H)**, and resting membrane potential **(I)**. P-values computed using Wilcoxon rank-sum test (*p<0.05, n=66 recordings for SAD B19, n=71 recordings for CVS-N2c).

We observed that samples infected with either CVS-N2c or SAD B19 RVdG showed an enrichment of neuronal populations, particularly L2/3 ExNs, over glial cells compared to uninfected cells (Figure 5B). This enrichment is likely due, in part, to the use of the human synapsin I promoter to drive helper virus transcripts, which biases starter cell infection toward neurons^59^, coupled with preferential spread of RVdG trans-synaptically between neurons^60,61^. Gene set enrichment analysis (GSEA) revealed an enrichment in inflammatory and immune response pathways in the SAD B19 datasets (Figures 5C-5D, S5). These signatures were primarily driven by a population of inflammatory astrocytes that were present only in the SAD B19 datasets (Supplement 5J and 5L), in line with previous studies^62–64^.

We also sought to assess strain-dependent toxicity using targeted whole-cell patch clamp recording of cells with neuronal morphology. SAD B19-infected cells showed blebbing of the axodendritic processes and were less likely to fire action potentials compared to CVS-N2c–infected cells (Figures 5E, 5F and S6). Intrinsic biophysical properties including cell capacitance, input resistance, and resting membrane potential were more variable in SAD B19–infected compared to CVS-N2c–infected neurons (Figure 5G–5I), and the resting membrane potential was significantly depolarized in SAD B19–infected compared to CVS-N2c–infected neurons (Figure 5I). Thus, while similar transcriptomic cell types are present in both the SAD B19–infected and CVS-N2c–infected cell populations, SAD B19-infected cells exhibit both gene expression electrophysiological signatures toxicity compared to CVS-N2c-infected cells.

### The MS2-MCP-KASH system can be used to target RVdG-encoded barcodes to the nuclear membrane for snRNA-seq readout

Prior versions of barcoded RVdG virus are compatible with scRNA-seq readout; however, adult brain tissue is more commonly studied using snRNA-seq due to loss of susceptible cell types such as projection neurons during single cell dissociation^58,65^. RVdG viral replication and transcription occurs entirely in the cytoplasm, preventing readout of RVdG-encoded barcodes from nuclei^66^. To overcome this limitation, we leveraged MS2 tagging, which takes advantage of strong interactions between MS2 bacteriophage coat protein (MCP) and a defined RNA sequence that forms a hairpin *in situ*^67,68^. First, we cloned an MCP tagged with the fluorescent protein oScarlet and the KASH outer nuclear membrane localization domain^69^ into the fourth open reading frame of the CVS N2c RVdG genome. We additionally cloned the 6x tandem MS2 hairpin sequence into the 3’ UTR of the MCP-oScarlet-KASH sequence proximal to the barcode sequence. Together, these two modifications provide a mechanism to target both the fluorescent reporter oScarlet and the RVdG-encoded barcode transcripts to the outer nuclear membrane, facilitating barcode recovery from sorted nuclei (Figures 6A–6B).

**Figure 6.**
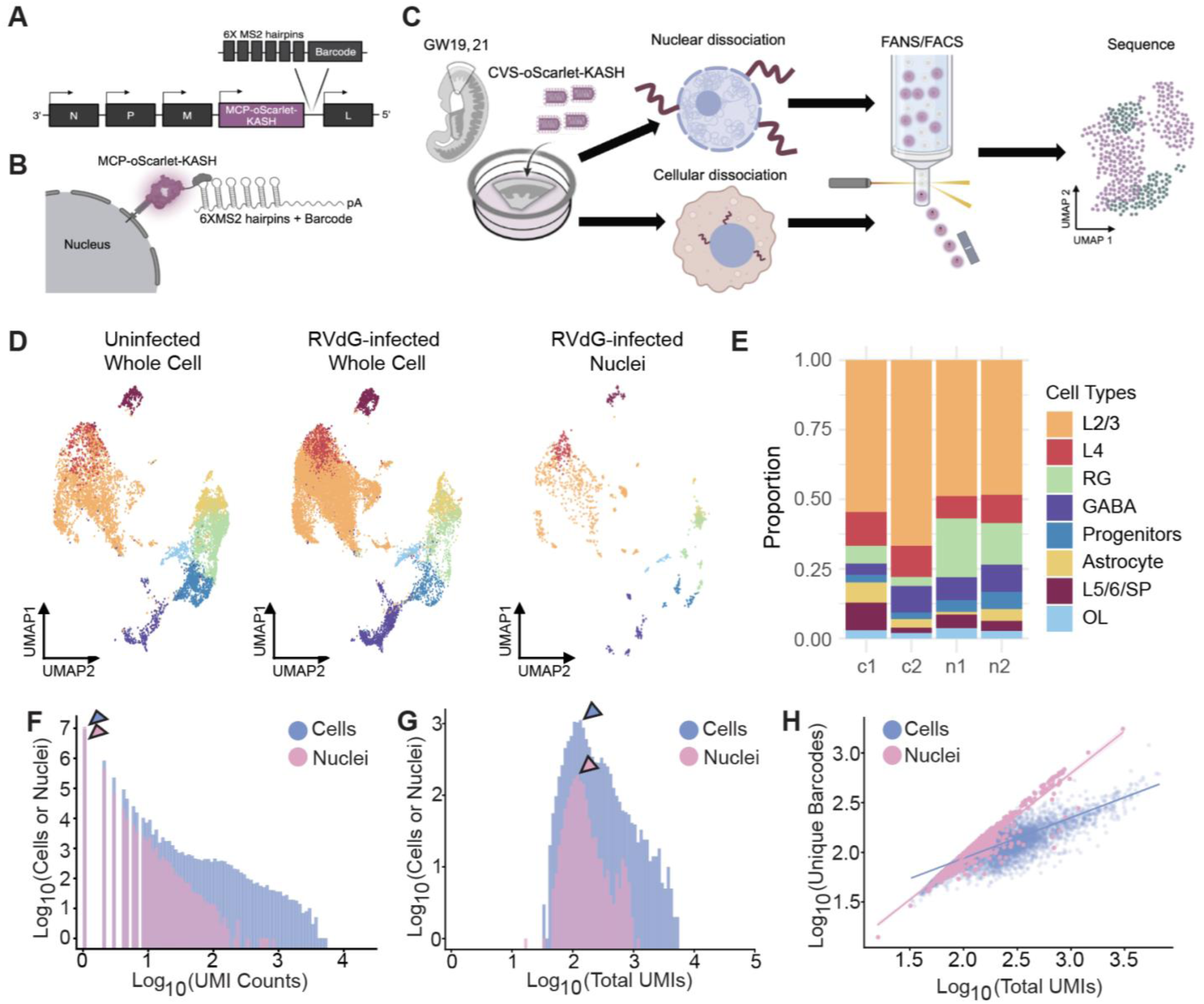
The MS2-MCP-KASH system can be used to target RVdG-encoded barcodes to the nuclear membrane for snRNA-seq readout. **A.** Location of barcode and nuclear localization signal cloning site in the rabies genome. **B.** Strategy for targeting of RVdG barcode transcripts to the nucleus utilizing KASH. **C.** Experimental paradigm in which slices from the same donors were dissociated into either single cells or single nuclei before enriching for RVdG-infected cells/nuclei for RNA-sequencing. **D.** UMAP visualizations of transcriptomic cell types from uninfected whole cell, RVdG-infected whole cell, and RVdG-infected nuclei datasets. **E.** Relative proportions of cell types captured by whole cell (c) and nuclear (n) datasets. **F.** Distribution of UMI counts for each unique barcode found within cells or nuclei. Medians are annotated by (*p<0.05). **G.** Distribution of total RVdG barcode UMIs captured per cell or nucleus. Medians are annotated by arrowheads (median=135 for cells and median=120 for nuclei). P-value computed using the Wilcoxon rank-sum test (*p<0.05). **H.** Correlation between total number of UMIs detected and number of unique barcodes detected per cell or nucleus with line of best fit. P-value computed using Spearman’s Correlation (cell R^2^=0.66, nuclei R^2^=0.90; *p<0.05).

To determine how well we could capture RVdG barcodes in nuclei using MS2-KASH tagging, in a subset of cases we isolated nuclei for snRNA-seq and whole cells for scRNA-seq in parallel from adjacent slices infected with the CVS-N2c-MS2-KASH-oScarlet RVdG (Figures 6C and S3C). Our snRNA-seq dataset included all of the major transcriptomic cell types we identified using scRNA-seq, indicating that nuclei isolations preserve the cellular diversity observed in whole cell dissociations (Figures 6D-6E). Within these paired samples, we identified 1,141 nuclei and 9,332 cells after filtering based on quality control metrics (STAR Methods). We performed targeted amplification of the RVdG barcodes (STAR Methods) and quantified RVdG barcodes after deduplication based on cell/nucleus barcode and UMI. In total, we detected 451,324 unique cell:barcode combinations and 115,635 unique nucleus:barcode combinations. We detected slightly fewer UMIs per unique cell/nucleus:barcode combination in nuclei compared to cells (Figure 6F; median=1 UMIs per unique cell:barcode combination, median=1 UMIs per unique nucleus:barcode combination) but similar numbers of total barcode UMIs in both (Figure 6G; median=135 total barcode UMIs per cell, median=120 total barcode UMIs per nucleus). There was a positive correlation between total barcode UMIs detected and number of unique barcodes detected in both cells and nuclei (Figure 6H). Taken together, these findings demonstrate that RVdG barcode capture from nuclei is tractable and only moderately less sensitive than barcode capture from cells. The ability to use snRNA-seq readout of RVdG barcodes provides a path forward for RV-based circuit mapping in more mature tissues that are not amenable to whole cell dissociation, such as the adult human brain.

### Reduction of ambient barcode sequences using UMI thresholding

Ambient RNA in a cell/nucleus suspension can be aberrantly counted along with a cell’s native mRNA during droplet-based sc/snRNA-seq, leading to cross-contamination of transcripts between different cell populations^70^. While low expression of ambient transcripts may be insufficient to alter the cell type classification of a sequenced cell or nucleus, it could potentially lead to false positive connections when reading out connectivity using RVdG-encoded barcodes. Therefore, we sought to establish quality control metrics for assigning RVdG barcodes to cells and nuclei while minimizing potential artifacts due to ambient RNA transcripts. In single cell datasets we detected RVdG barcodes in 5.85% of empty droplets, defined as containing ≤10 total UMIs in the paired cell transcriptome library (Figure 7A). Empty droplets contained fewer total barcode UMIs (median=136 in real cells, median=1 in empty droplets; Figure 7B) and fewer unique barcodes compared to real cells (median=99 in real cells, median=1 in empty droplets; Figures 7C and 7D). To minimize the risk of inferring false positive connections due to ambient barcodes, we aimed to identify a UMI threshold to remove ambient barcodes from downstream analysis (Figure 7E). We found that implementing a threshold of ≥3 UMI counts per cell:barcode combination was sufficient to filter out more than 90% of empty droplets while retaining more than 90% of cells (Figure 7E). We performed a similar analysis on our datasets generated from single nuclei dissociations which revealed RVdG barcodes in 5.06% of empty droplets (Figure 7F). Real nuclei also had more total barcode UMIs (median=116 in real nuclei, median=1 for empty droplets) and more unique barcodes (median=100 in real nuclei, median=1 in empty droplets) compared to empty droplets (Figure 7G, 7H, and 7I).

**Figure 7.**
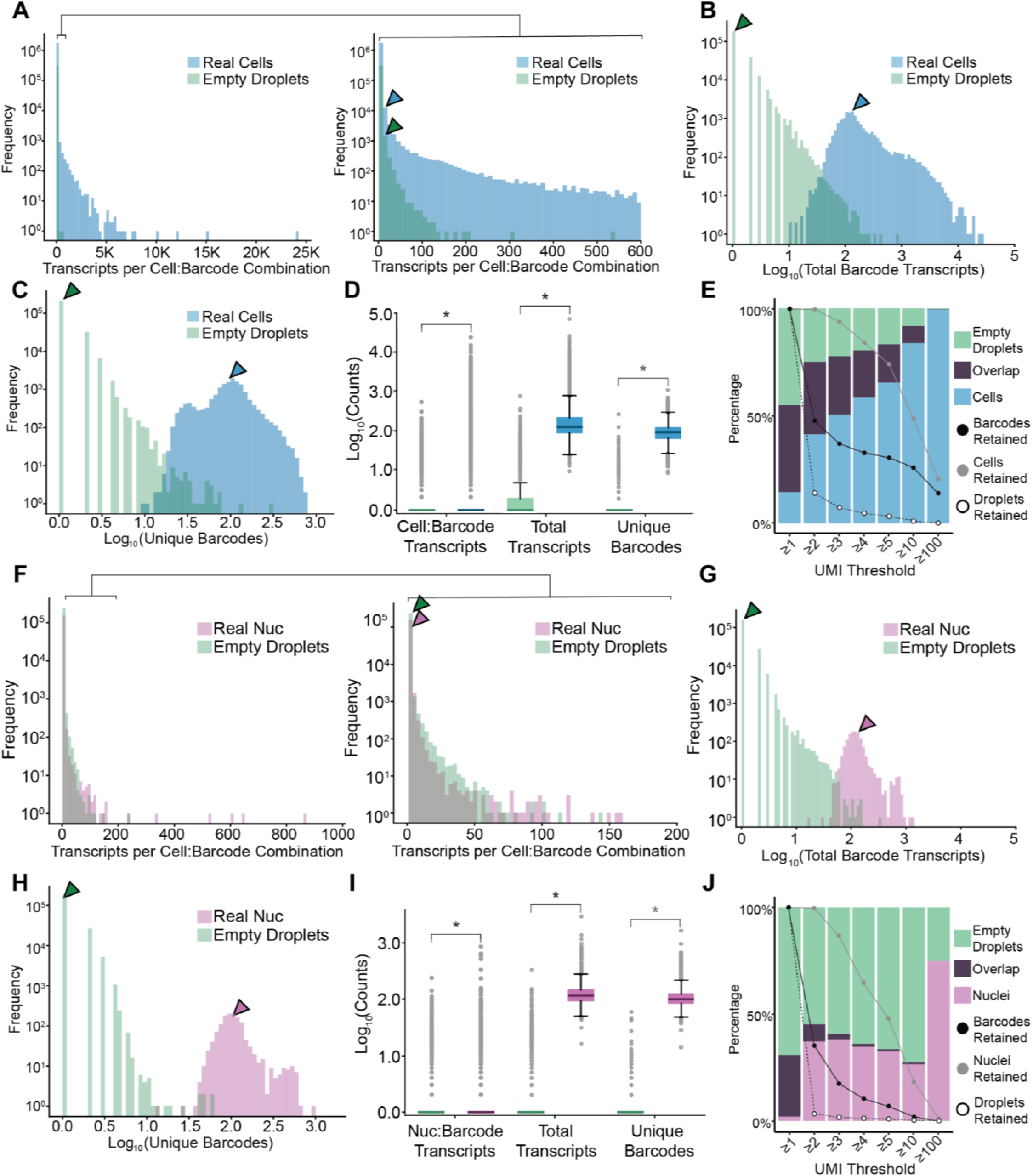
Reduction of ambient barcode sequences using UMI thresholding. **A.** Number of UMI transcripts detected for every barcode in each cell or empty droplet from single cell dissociation experiments, with magnified panel showing distribution from 0-600 UMIs on right (median=1 in both real cells and empty droplets, indicated by arrowheads; n=1,740,015 cell:barcode combinations, n=315,450 empty droplet:barcode combinations). **B.** Distribution of total barcode UMI transcripts detected per cell or empty droplet (median=136 in real cells, median=1 in empty droplets, indicated by arrowheads; n=16,036 cells, n=253,000 empty droplets). **C.** Number of unique barcodes detected per cell or empty droplet (median=99 in real cells, median=1 in empty droplets, indicated by arrowheads; n=16,036 cells, n=253,000 empty droplets). **D.** Number of UMI transcripts per each barcode, total barcode UMI transcripts, and number of unique barcodes per cell or empty droplet. Box plot shows median and interquartile range (IQR) of data in **(A–C)**. **E.** Stacked bar plot showing fraction of barcodes uniquely found in empty droplets, cells, or both groups after implementing the indicated UMI threshold. Percent of RVdG barcodes, cells, and empty droplets retained after implementing the UMI threshold is overlaid. **F.** Number of UMI transcripts detected for every barcode in each nucleus or empty droplet from single nucleus dissociation experiments, with magnified panel showing distribution from 0-200 UMIs on right (median=1 in both real nuclei and empty droplets, indicated by arrowheads; n=157,073 nuclei:barcode combinations, n=239,338 empty droplet:barcode combinations). **G.** Distribution of total barcode UMI transcripts detected per nucleus or empty droplet (median=116 in real nuclei, median=1 in empty droplets, indicated by arrowheads; n=1,307 nuclei, n=196,934 empty droplets). **H.** Number of unique barcodes detected per nucleus or empty droplet (median=100 in real nuclei, median=1 in empty droplets; n=1,307 nuclei, n=196,934 empty droplets). **I.** Number of UMI transcripts per each barcode, total barcode UMI transcripts, and unique barcodes per nucleus or empty droplet. Box plot shows median and IQR of data in **(F–H)**. **J.** Stacked bar plot showing fraction of barcodes uniquely found in empty droplets, nuclei, or both groups after implementing the indicated UMI threshold. Percent of RVdG barcodes, nuclei, and empty droplets retained after implementing the UMI threshold is overlaid. In **(D)** and **(I)**, p-values are computed using the Wilcoxon rank-sum test, *p<0.05.

Applying the threshold to keep only nucleus:barcode combinations with a UMI count ≥3 filtered out approximately 98% of empty droplets while retaining 87% of nuclei for downstream analysis (Figure 7J). Based on these results, only cell/nucleus:barcode combinations with ≥3 UMIs were used for downstream connectivity analysis.

### Barcoded RVdG circuit mapping reveals the emergence of cell type–specific connectivity motifs in second trimester human cerebral cortex

To assess the emerging connectivity matrix in second trimester human cerebral cortex, we pooled together our single-cell and single-nuclei sequencing datasets and their corresponding RVdG barcodes (Table S4). We identified a total of 12,036 unique RVdG barcodes (Figure S7A). To identify putative connections we identified networks of cells that shared a common barcode (‘barcode networks’), and annotated cells as starter or non-starter cells based on expression of helper virus (STAR Methods). Barcode networks with no starter cell were excluded from further analysis. We identified 5,047 barcodes in 1,230 starter cells: 2,762 barcodes were found in a single starter cell and 2,285 barcodes were found in multiple starter cells (Figure 8B). The barcode networks containing multiple starter cells were excluded from further analysis due to the ambiguity of connection directionality and possibility of polysynaptic spread through multiple starter cells. Of the 2,762 barcodes found in a single starter cell, 1,102 were also found in at least one non-starter cell. We proceeded to infer connectivity using only the 457 barcode networks containing a single starter cell and at least one non-starter cell (‘single starter networks’), where directionality can be determined based on retrograde spread of RVdG virus^1^. To prevent calling duplicate connections when multiple barcodes are shared between two cells, we collapsed all instances of connections between the same two cells into a single count. Distinct barcode networks with converging cells were then collapsed into singular networks, and duplicate connections were merged into one labeled connection (Figure 8A). After collapsing all duplicate connections, we identified 457 singular networks consisting of 4,331 directed connections centered around individual starter cells (median=5 cells per network, Figure 8C). The cell type composition in each network was similar across different network sizes (Figure 8D-E) and the vast majority of connections (3622/4331, 83.6%) were between neuronal cell types.

**Figure 8.**
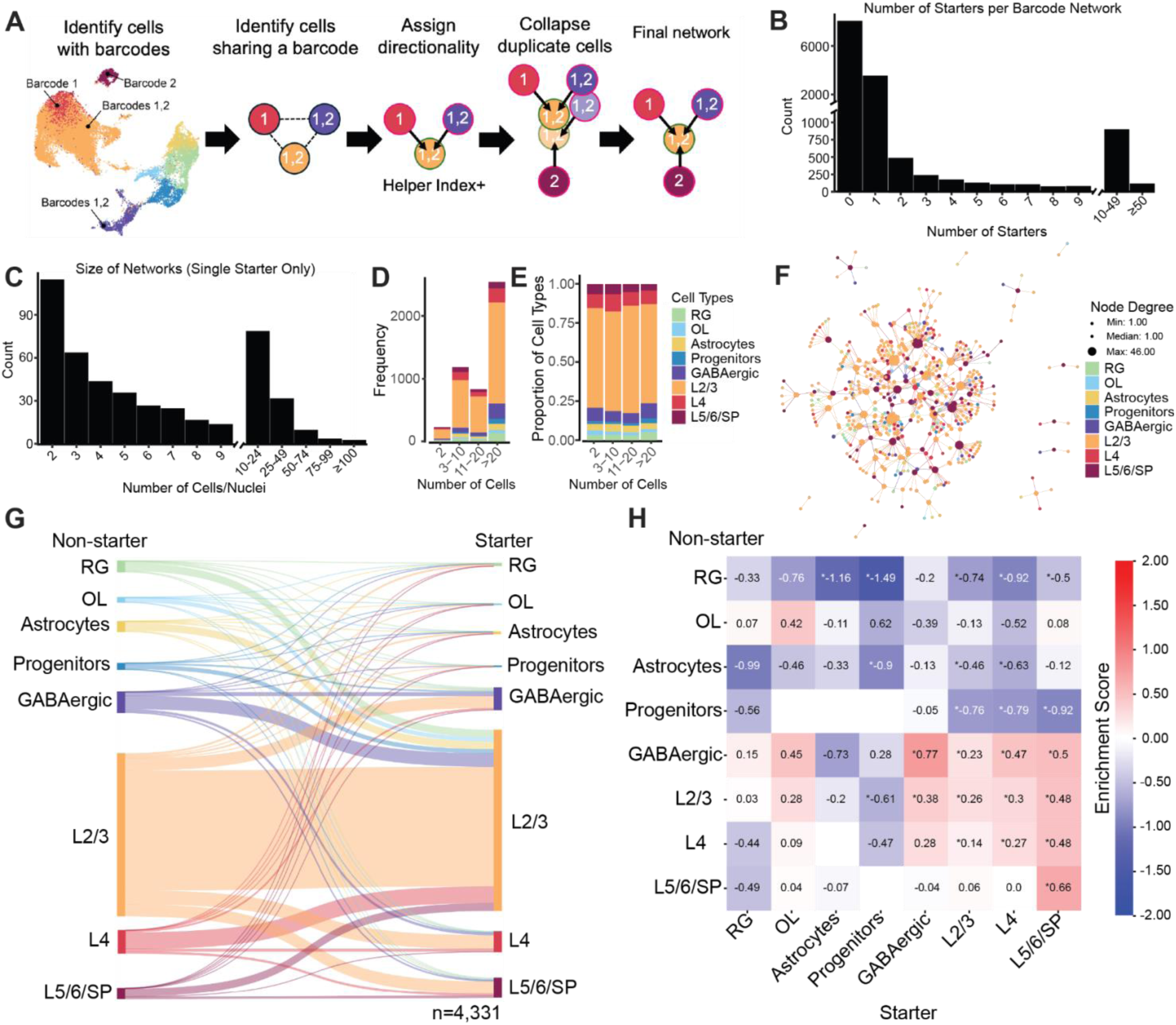
Barcoded RVdG circuit mapping reveals the emergence of cell type–specific connectivity motifs in second trimester human cerebral cortex. **A.** Schematic of connectivity analysis. **B.** Number of starter cells per barcode network. **C.** Distribution of total number of cells per single-starter barcode network. **D.** Frequency of barcode networks containing different numbers of cells. Cell type proportions for each network size interval are indicated by color. **E.** Proportions of cell types in different barcode network size intervals. **F.** Example graph network showing the merged networks in one tissue slice infected with CVS-N2c RVdG. **G.** Sankey diagram showing observed connections by cell type (n=4,331 connections). **H.** Normalized connectivity matrix showing enrichment scores of observed connections compared to a random connectivity matrix weighted by starter and uninfected cell type proportions (STAR Methods). P-values are computed using the two-sided binomial tests with Bonferroni correction for multiple comparisons; *p<0.05.

Next we merged singular networks together to visualize them as graphs in which each cell appears only once, by linking non-starter cells that could exist in multiple singular networks (Figures 8F and S8). A plurality of connections were between L2/3 ExNs (n=2039/4331, 47.08%) and L2/3 neurons received the largest proportion of inputs compared to other cell types (n=3087/4331, 71.28%; Figure 8G, S7B). Given that L2/3 ExNs are also the most prevalent cell type in our dataset, irrespective of RVdG infection (Figures 4G and 5B), we next asked how the observed connectivity matrix compared to a null connectivity matrix with random connections between the defined starter cell population and the pool of possible interacting cells. Specifically, the connectivity expected by chance between any two cell types was modeled as a binomial distribution considering the proportions of those cell types in the starter and uninfected populations (STAR Methods). After controlling for cell type proportions, neuronal cell types were still enriched in connections and glial cell types were depleted of connections (Figure 8H), consistent with RVdG spread being largely restricted to chemical synapses in the developing human brain. The strongest enrichment was observed among GABAergic neurons, which also target all other neuronal cell types more often than predicted by chance. In contrast, L5/6/SP receive inputs from all neuronal populations, but preferentially target other L5/6/SP neurons. Taken together, these findings suggest that cell type–specific circuit motifs are already beginning to emerge in midtrimester human brain development.

## DISCUSSION

Understanding neuronal connectivity at the level of molecularly defined cell types remains a key challenge in neuroscience. Sequencing-based approaches to connectivity mapping provide a potential path forward for scalable circuit mapping by leveraging the power and throughput of single cell genomics. In particular, barcoded RVdG has recently emerged as one strategy for integrated circuit mapping and molecular interrogation of the mapped cells^3–5^, but its application has been limited by the degree of barcode complexity and an inability to reliably detect RVdG transcripts in dissociated single nuclei^71^. Here we introduce an RVdG packaging protocol that is specifically tailored to improve barcode complexity, increasing the number of cells that can be mapped simultaneously in one experiment by several orders of magnitude compared to prior protocols. We also describe a strategy for nuclear localization of RVdG barcodes that enables snRNA-seq readout of the RVdG-encoded barcodes. Lastly, we demonstrate the power of these new tools by mapping local cell type–specific connectivity in organotypic slice cultures of the developing human cerebral cortex. By converting the problem of synaptic connectivity into a problem of barcode sequencing, barcoded RVdG circuit mapping has the potential to transform connectomics into a routine experiment that can be performed in any lab with modest resources, making it possible to explore how circuits differ between treatment conditions, in disease states, between the sexes, across the lifespan and across species.

Barcode complexity is a limiting feature of all barcoded viral tracing strategies^72^, but has been especially problematic for barcoded RVdG circuit mapping due to the low efficiency of viral barcode recovery during the packaging process^3^. We identified two key factors that improve the complexity of barcoded RVdG libraries: transfection reagent and packaging cell line. Specifically, of the options we tested, Lipo3000 and PEI MAX offer a substantial improvement in the percentage of cells recovering active RVdG particles compared to other reagents including Lipo2000^3,5^ or Xfect^4^. A direct comparison of library complexity between RVdG packaged in Lipo3000 and Lipo2000 revealed a significant improvement in barcode complexity in libraries that were packaged in Lipo3000, suggesting that improved transfection efficiency directly translates to improved barcode complexity. We also found that HEK293T^4^ is more efficient at rescuing RVdG compared to the BHK21-derived B7GG fibroblast line^3^ and Neuro2A^5^, in line with previous work^6,73^. One potential argument for packaging RVdG in Neuro2A cells is that it may enhance neurotropism and decrease neurotoxicity^58^. However, at least one recent study found no significant difference in connectivity data acquired using RVdG recovered in HEK293T cells or Neuro2A cells^17^. In addition, we show that a HEK293T cell line that stably expresses oG (HEK-GT^17^) further improves uniformity of barcode libraries. There could be several reasons for this. First, the dynamic range of rabies G expression is likely narrower in the cell line compared to plasmid-based transient transfection^74^. Because rabies G confers expression-level dependent spread^18,75^, this may provide a more fair competition between expanding barcode clones. Additionally, oG shows up to 20-fold greater efficiency in promoting transsynaptic spread^18^ compared to SAD B19 G, which is traditionally used for packaging RVdG. Finally, transfecting fewer plasmids may improve overall transfection efficiency^76^. New transfection reagents, additives, and cell lines that stably expresses the additional genes required for RVdG packaging may further enhance barcoded RVdG library complexity.

We used a defined barcode whitelist^11^ in lieu of randomer^3,5^ or semi-randomer^4^ barcodes which can require complex probabilistic models to determine whether similar but non-identical barcodes represented the same or distinct sequences. Using a pre-defined whitelist of unique sequences allows us to ensure that any two barcodes differ by at least 6 bp (Hamming distance ≥ 6), making barcode identification robust to sequencing errors or mutations that arise during virus production. Furthermore, by using the whitelist as a reference genome, barcodes can be aligned using standard transcriptomic aligners (e.g., STAR, bowtie2), streamlining the process and reducing the computational complexity typically required to collapse similar barcode sequences^4^. However, our barcodes are 61 bp in length, requiring longer sequencing reads that may preclude spatial transcriptomic readout used by several groups for barcoded RVdG circuit mapping^5,77^. New barcode strategies that can balance a high Hamming distance with short overall barcode length might further streamline the barcode design to such that it can be read out using any platform.

Rabies virus is an RNA virus that is localized primarily within the cytoplasm^9,10^ and prior studies utilized whole-cell single-cell transcriptomics for barcode recovery. However, more mature brain tissue and flash frozen tissue cannot be easily dissociated into intact cells. We developed a method for localizing barcoded transcripts to the nucleus by leveraging MS2 tagging^67,68^ guided by the KASH nuclear localization signal, and demonstrated efficient recovery of RVdG-encoded barcode transcripts from single nuclei. It is possible that tethering the barcode transcripts to a different nuclear protein, such as histone 2B (H2B), could further improve barcode recovery in nuclei.

We focused on two commonly used RVdG strains: SAD B19 and CVS-N2c. Consistent with prior reports, our data suggest that the CVS-N2c strain is less toxic than the SAD B19 strain^58^. Despite this impact on cell health, we were able to generate high-complexity libraries using both strains. Circuit mapping experiments showed that both strains captured similar proportions of cell types. Further experiments aimed at elucidating nuanced differences in the tropism of different RVdG strains, and the rabies G that drives trans-synaptic spread, will be important to be able to compare RVdG-inferred connectomes to each other and to ground truth wiring diagram of a circuit, which may be difficult to know a priori.

We applied barcoded RVdG to explore connectivity in organotypic slice cultures of the human brain during the second trimester of gestation, a developmental window during which neuronal connectivity is actively being established. We observed an enrichment in connectivity between neuronal cell types compared to non-neuronal cell types, supporting the idea that rabies virus is biased towards trans-synaptic spread. We observed the greatest enrichment in connectivity between GABAergic neurons, and from GABAergic to other neuronal populations. Previous studies have demonstrated the critical role of depolarizing GABAergic signaling in the formation of AMPA receptor-mediated glutamatergic circuitry^78–81^. This mechanism is thought to be mediated by synaptic release of GABA onto maturing glutamatergic neuronal populations, and our data suggest that this may be one of the most prominent circuit motifs during mid-gestation in humans. We also observed enrichment in connectivity from all neuronal populations onto L5/6/SP ExNs, consistent with previous work^78,82–84^. In particular, the enrichment in connections from L4 to L5/6/SP may correspond to transient synapses formed during the establishment of the thalamus-subplate-thalamus circuit. We also identify shared barcodes between neurons and non-neuronal cell types, suggesting RVdG spread is not entirely restricted to mature synapses^3,60,61,85^. Shared barcodes between cells may also arise following division of a single infected cell, such as an IPC, RG, or oligodendrocyte precursor cell (OPC, a subset of OL in our dataset). Disentangling the contribution of cell division and trans-synaptic spread in actively dividing cell populations may be possible in future studies by using two barcode systems in parallel - one for tracking cell lineage (e.g. a barcoded lentivirus) and another for tracking connectivity (barcoded RVdG).

In summary, we report an optimized protocol for generation of high-complexity barcoded RVdG that is compatible with both single-cell and single-nucleus RNA-sequencing readout. This powerful new tool enables probing synaptic connectivity at unprecedented scale and molecular detail. Future applications of this tool may help to reveal the organizational principles of brain circuits across species, ages, and disease states.

## ACKNOWLEDGEMENTS

We thank members of the Nowakowski and Cadwell labs for providing helpful feedback throughout this project and Ken Probst and Melissa Logies for assistance with figure layout and schematics. This work was supported, in part, as grants from NIH: U01MH130700, R01NS123263, R01MH128364 and RF1MH125516 (to T.J.N.), K08NS126573 and U01NS132353 (to C.R.C), Awards from the Klingenstein-Simons Fund in Neuroscience, Broad Foundation and Sontag Foundation (to T.J.N.), the Shurl and Kay Curci Foundation (to T.J.N. and C.R.C.), and the Weill Neurohub (to C.R.C.), and gifts from Schmidt Futures and the William K. Bowes Jr Foundation (to T.J.N). This work was also supported by the National Science Foundation under award number NSF #2134955 (T.N). T.J.N. is a New York Stem Cell Foundation Robertson Neuroscience Investigator. D.S. was supported by the NSF Graduate Research Fellowship Program (GRFP). M.E.U was supported by the NSF GRFP and the Hertz Foundation Fellowship. Sequencing was performed at the UCSF CAT, supported by UCSF PBBR, RRP IMIA, and NIH 1S10OD028511 grants.

## AUTHOR CONTRIBUTIONS

D.S., D.B., T.J.N., and C.R.C. conceptualized the study. D.S., H.H.L., M.E.U., K.L., D.L.B., and E.S.-B. performed experiments. M.E.U., A.J.M., D.S., S.R. and C.R.C. performed analyses. D.S., M.E.U., A.J.M. and C.R.C wrote and edited the manuscript with input from all other authors. C.R.C and T.J.N. supervised the study and acquired funding to support this work.

## DECLARATION OF GENERATIVE AI AND AI-ASSISTED TECHNOLOGIES

During the preparation of this work, the authors used ChatGPT to suggest changes to improve the clarity of the text. After using this tool or service, the authors reviewed and edited the content as needed and take full responsibility for the content of the publication.

## STAR★METHODS

### KEY RESOURCE TABLE

| REAGENT or RESOURCE | SOURCE | IDENTIFIER |
| --- | --- | --- |
| **Antibodies** |  |  |
| NeuN (chicken, 1:500) | MilliporeSigma | ABN91 |
| NeuN (guinea pig, 1:500) | MilliporeSigma | ABN90 |
| VGLUT1 (mouse, 1:200) | MilliporeSigma | MAB5502 |
| PSD-95 (rabbit, 1:350) | Invitrogen | 51-6900 |
| SYP (guinea pig, 1:1000) | Synaptic Systems | 101004 |
| GFP (chicken, 1:1000) | Aves | 1020 |
| dsRed (rabbit, 1:250) | Takara | 632496 |
| RFP (rat, 1:250) | Chromotek | 5F8 |
| DLX2 (mouse, 1:50) | Santa Cruz | sc-393879 |
| NEUROD2 (rabbit, 1:250) | Abcam | ab104430 |
| **Bacterial and Viral Strains** |  |  |
| 5-alpha Competent E. Coli | New England BioLabs | C2987H |
| NEB Stable Competent E. Coli | New England Biolabs | C3040H |
| Stellar™ Competent Cells | Clontech Labs | 636763 |
| MegaX DH10B T1<sup>R</sup> Electrocomp™ Cells | Thermo Fisher | C640003 |
| **Chemicals, Peptides, and Recombinant Proteins** |  |  |
| Heat-inactivated FBS | HyClone | SH30071.03HI |
| DMEM with Glutamax | Gibco | 10569044 |
| 0.25% Trypsin-EDTA | Gibco | 25200114 |
| Poly-D-Lysine | MilliporeSigma | P7280 |
| Polybrene | MilliporeSigma | TR-1003-G |
| Trans-Isrib | Tocris | 5284 |
| Chroman 1 | Medchem Express | HY-15392 |
| Emricasan | Selleckchem | S7775 |
| Polyamine Supplement | Sigma-Aldrich | P8483 |
| Glutamax | Gibco | 35050-061 |
| N2 supplement | Gibco | 17502-048 |
| B27 w/ Vitamin A | Gibco | 17504-001 |
| Neurobasal | Gibco | 21103049 |
| DMEM/F12 with Glutamax | Gibco | 10565042 |
| L-Ascorbic Acid 2-Phosphate | Wako Chemicals USA | 321-44823 |
| Non-essential amino acids | Gibco | 11140050 |
| Primocin | Fisher Scientific | NC9141851 |
| Donkey Serum | Jackson Immuno | 017-000-121 |
| Prolong Gold | ThermoFisher | P36934 |
| Fluoromount-G | ThermoFisher | 00-4959-52 |
| PEG8000 | Promega | V3011 |
| Sodium Chloride | Spectrum Chemical | SO160 |
| 1X PBS | Gibco | 10010023 |
| Trizol | ThermoFisher | 15596026 |
| Chloroform | Fisher Scientific | 60-047-878 |
| Phenol:Chloroform:Isoamyl Alcohol<br>(25:24:1, v/v) | ThermoFisher | 15593031 |
| Glycoblue | ThermoFisher | AM9515 |
| Ethanol, 200 proof | Weill Chem Central | TX89125-172SFU |
| Isopropyl Alcohol | Millipore Sigma | 563935 |
| Sodium Acetate, 3M | ThermoFisher | J61928.AE |
| Lipofectamine 3000 | ThermoFisher | L3000015 |
| Lipofectamine 2000 | ThermoFisher | 11668019 |
| XFect | Takara | 631318 |
| XtremeGene HP | Roche | 06366236001 |
| FuGENE HD | Promega | E2311 |
| PEI MAX | PolySciences | 49553-93-7 |
| TransIT-293 | Mirus Bio | MIR2704 |
| Q5 HotStart 2X Master Mix | New England BioLabs | M0494L |
| KAPA HotStart ReadyMix 2X | Fisher Scientific | NC0295239 |
| EBSS | Worthington | LK003188 |
| DNase I | MilliporeSigma | DN25-1 G |
| Trehalose | MilliporeSigma | T9531-25G |
| Potassium Chloride | MilliporeSigma | 529552-1KG |
| Sodium Phosphate Monobasic | MilliporeSigma | S0751-1KG |
| Sodium Bicarbonate | MilliporeSigma | S5761-1KG |
| Glucose | MilliporeSigma | G7021-1KG |
| Magnesium Chloride Solution | MilliporeSigma | M1028-100ML |
| Calcium Chloride Solution | MilliporeSigma | 21115-100ML |
| Potassium Gluconate | MilliporeSigma | P1847-100G |
| HEPES Solution | MilliporeSigma | H0887-100ML |
| Adenosine 5'-triphosphate Magnesium Salt | MilliporeSigma | A9187-100MG |
| Guanosine 5'-triphosphate Sodium Salt | MilliporeSigma | G8877-10MG |
| Sodium Phosphocreatine | MilliporeSigma | P7936-1G |
| Biocytin | MilliporeSigma | B4261-50MG |
| Potassium Hydroxide Solution | MilliporeSigma | P4494-50ML |
| EGTA | MilliporeSigma | E3889-100G |
| Recombinant RNase Inhibitor | Roche | 09537589103 |
| **Critical Commercial Assays** |  |  |
| PIPseq T2 v4.0PLUS | Fluent Biosciences | FBS-SCR-T2-8-V4.05-FLU |

| **Deposited Data** |  |  |
| --- | --- | --- |
| scRNA-seq/snRNA-seq datasets | Raw: dbGAP<br>Processed: UCSC Cell Browser | Raw: In Submission<br>Processed: https://barcoded-rabies-cortex.cells.ucsc.edu |
| Barcoded RVdG Genome Library Sequencing Datasets | GEO | GSE277536 |
| **Experimental Models: Cell Lines** |  |  |
| LentiX 293T | Takara | 632180 |
| HEK-GT | Ben-Simon Lab | https://elifesciences.org/articles/79848 |
| BHK-EnvA-TVA | Ben-Simon Lab | https://elifesciences.org/articles/79848 |
| BHK-EnvA | Callaway Lab | Salk Institute Virus Core: BHK-EnvA |
| Neuro2A | ATCC | CCL-131 |
| 293T-TVA800 | Callaway Lab | Salk Institute Virus Core: 293T-TVA800 |
| **Oligonucleotides: see Table S2** |  |  |
| **Recombinant DNA** |  |  |
| pMD2.G | Addgene | 12259 |
| psPAX2 | Addgene | 12260 |
| pcDNA 3.1 puro Nodamura B2 | Addgene | 17228 |
| Barcoded SAD B19 RVdG dTomato genome plasmid | This Study | Will be submitted to Addgene |
| Barcoded CVS-N2c RVdG MCP- oScarlet-KASH genome plasmid | This Study | Will be submitted to Addgene |
| pCAG-B19N | Addgene | 59924 |
| pCAG-B19P | Addgene | 59925 |
| pCAG-B19G | Addgene | 59921 |
| pCAG-B19L | This Study | 59922 |
| pCAGGS-T7opt | This Study | 65974 |
| pSicoFlipped-syn-GFP-TVA-oG-WPRE-bGHpA | This Study | Will be submitted to Addgene |
| pSicoFlipped-syn-GFP-TVA-WPRE-bGHpA | This Study | Will be submitted to Addgene |
| pSicoFlipped-syn-GFP-TVA-oG-6xMS2-WPRE-bGHpA | This Study | Will be submitted to Addgene |
| **Software and Algorithms** |  |  |
| ImageJ (Fiji) | ImageJ | https://imagej.net/Fiji |
| pClamp v11 | Molecular Devices | https://www.moleculardevices.com/products/axon-patch-clamp-system/acquisition-and-analysis-software/pclamp-software-suite |
| PipSeeker v3.3 | Fluent Biosciences | https://www.fluentbio.com/products/pipseeker-software-for-data-analysis/ |
| R v4.3.3 | R | https://www.r-project.org/ |
| Python v3.11.4 | Python | https://www.python.org/ |
| JupyterLab v4.0.2 | Project Jupyter | https://jupyter.org/ |
| RStudio v2023.06.0+421 | Posit | https://posit.co/download/rstudio-desktop/ |
| Seurat v5.0 | Satija Lab | https://satijalab.org/seurat/ |
| Devtools v2.4.5 | RStudio | https://devtools.r-lib.org/ |
| BiocManager v1.30.23 | Martin Morgan | https://bioconductor.github.io/BiocManager/ |
| SeuratObject v5.0.2 | Satija Lab | https://github.com/satijalab/seurat-object |
| dplyr v1.1.4 | TidyVerse | https://dplyr.tidyverse.org |
| Ggplot2 v3.5.1 | TidyVerse | https://ggplot2.tidyverse.org |
| EnhancedVolcano v1.20.0 | Kevin Blighe | https://github.com/kevinblighe/EnhancedVolcano |
| harmony v1.2.0 | Raychaudhuri Lab | https://github.com/immunogenomics/harmony |
| UCell v2.6.2 | Carmona Lab | https://github.com/carmonalab/UCell |
| clusterProfiler v4.10.1 | Yu Lab | https://github.com/YuLab-SMU/clusterProfiler |
| Msigdbr v7.5.1 | Igor Dolgalev | https://igordot.github.io/msigdbr/ |
| pandas v2.1.4 | pandas Development Team | https://pandas.pydata.org/ |
| NumPy v1.26.2 | NumPy | https://numpy.org/ |
| Matplotlib v3.8.2 | Matplotlib Development Team | https://matplotlib.org/ |
| SciPy v1.11.1 | SciPy | https://scipy.org/ |
| seaborn v0.12.2 | Michael Waskom | https://seaborn.pydata.org/ |
| statsmodels v0.14.3 | statsmodels | https://www.statsmodels.org/stable/index.html |
| NetworkX v3.3 | NetworkX Developers | https://networkx.org/ |
| graphviz v0.20.3 | graphviz Developers | https://pypi.org/project/graphviz/#description |
| SankeyMATIC | Steve Bogart | https://sankeymatic.com/build/ |
| Affinity Designer | Affinity | https://affinity.serif.com/en-us/designer/ |
| BioRender | BioRender | https://biorender.com/ |
| Bowtie2 v2.5.1 | Bowtie2 | https://bowtie-bio.sourceforge.net/bowtie2/index.shtml |
| STAR v2.5.3a | https://sourceforge.net/projects/bbmap/ | https://github.com/alexdobin/STAR/ |
| BBMap v39.01 | JGI BBTools | https://sourceforge.net/projects/bbmap/ |
| UMI-tools v1.1.4 | CGAT | https://github.com/CGATOxford/UMI-tools |
| Easy Electrophysiology | Easy Electrophysiology Ltd. | https://www.easyelectrophysiology.com/ |
| **Other** |  |  |
| Cell Culture Inserts, 0.4 µm pore size | MilliporeSigma | PICM03050 |
| Dounce | Wheaton | EW-04468-00 |
| Leica Widefield-Thunder Imaging System | Leica | UCSF Weill Microscopy Core |
| Leica SP8 Confocal Microscope | Leica | Nowakowski Lab |
| NextSeq 2000 | Illumina | Canzio Lab, UCSF |
| NovaSeq X | Illumina | UCSF Center for Advanced Technology |
| FACSAria Fusion | BD Biosciences | UCSF Laboratory for Cell Analysis |
| BX51WI Microscope | Olympus | BX51WI |
| 6-8 MΩ Resistance Glass Microelectrodes | Sutter Instruments | BF150-86-10 |
| Multiclamp 700B Amplifier | Molecular Devices | 700B |
| Digidata 1550B | Molecular Devices | 1550B |

### RESOURCE AVAILABILITY

#### Materials Availability

Plasmids generated in this study will be made available through Addgene. Other reagents are available from the Lead Contact, Dr. Tomasz Nowakowski, upon reasonable request.

#### Data and Code Availability

scRNA-seq and snRNA-seq raw data will be made available through dbGAP, and processed data are available through the UCSC Cell Browser: https://barcoded-rabies-cortex.cells.ucsc.edu. Sequencing data from barcoded viral genome and plasmid libraries for assessing diversity is available through GEO (GSE277536, private token: craxaqkirzkrtit). Other data that support the findings of this study are available from the Lead Contact, Dr. Tomasz Nowakowski upon reasonable request. All code used for analysis has been deposited at Github and will be publicly available as of the date of publication.

## EXPERIMENTAL MODEL AND SUBJECT DETAILS

### Human Primary Cerebral Cortex Specimens

De-identified tissue samples were collected with previous patient consent in strict observance of the legal and institutional ethical regulations. Protocols were approved by the Human Gamete, Embryo, and Stem Cell Research Committee (Institutional Review Board) at the University of California, San Francisco.

## METHOD DETAILS

### Generation of lentiviral helper plasmid and barcoded RVdG plasmid library

We cloned a lentiviral helper plasmid in which the human synapsin I promoter (hSyn1) drove the expression of a human codon–optimized GFP, TVA, and chimeric glycoprotein oG^18^ (pSico-hSyn1::GFP-T2A-TVA-E2A-oG). A modified pSico plasmid^11^ was utilized as the backbone, and the sequence between the cPPT/CTS and 3’ LTR was replaced with DNA fragments containing the hSyn1 promoter and human codon–optimized sequences GFP, T2A, TVA, E2A, and oG via gibson assembly using HiFi DNA Assembly Master Mix (NEB, E2621).

To prepare a SAD B19 strain rabies genome plasmid for barcoding (pSADdeltaG-dTomato), the R0558) to replace the gene encoding tagBFP2 with a gblock (IDT) containing a gene encoding a human codon–optimized dTomato and a barcode cloning site in the 3’ UTR of dTomato represented by two SacII restriction enzyme sites. To generate the backbone for the barcoded plasmid library, 10 μg of pSADdeltaG-dTomato was digested using SacII (NEB, R0157) overnight and purified using the Zymo DNA Clean and Concentrator-5 kit (Zymo, D4004). The insert containing the barcode library was generated by performing PCR on the STICR barcoded lentiviral plasmid library^11^ (Addgene 180483) using primers flanking the barcode sequences and containing overhangs matching the SacII-digested pSADdeltaG-dTomato plasmid (Primers 1 and 2, Table S2).

Similarly, to prepare a CVS-N2c strain rabies genome plasmid for barcoding and nuclear capture, the RabV CVS-N2c(deltaG)-mCherry plasmid (Addgene #73464) was digested with SmaI (NEB, R0141) and NheII-HF (NEB, R3131) to replace the gene encoding mCherry with a gblock (IDT) encoding a fusion protein consisting of the MS2 coat protein MCP, the fluorescent protein oScarlet, and the KASH nuclear localization signal. The 3’ UTR of this gblock carried six MS2 hairpin sequences followed by a barcode cloning site represented by two SacII restriction enzyme sites. 10 μg of the resulting plasmid was digested using SacII (NEB, R0157) overnight and purified using the Zymo DNA Clean and Concentrator-5 kit (Zymo, D4004). The insert containing the barcode library was generated by performing PCR on the STICR barcoded lentiviral plasmid library^11^ (Addgene 180483), with overhangs matching the SacII digested plasmid (Primers 1 and 2, Table S2).

Based on our initial qPCR optimization (Figure S1C), 10 cycles of PCR were performed on 25 ng of plasmid DNA in 50 μL total volume using KAPA HotStart PCR ReadyMix (Roche KK2601), with the following parameters: Initial denaturation: 95C for 3 min, Denaturation: 98C for 20 s, Annealing: 62C for 15 s, Extension: 72C for 10 s. A total of 5 PCR reactions was performed, which represents a starting amount of approximately 1.3×10^10^ molecules of DNA, or ∼100-fold excess of the theoretical diversity of the plasmid library. Each PCR reaction was subsequently supplemented with 5 μL of CutSmart Buffer (NEB B7204) and digested using 1μL DpnI restriction enzyme (NEB R0176) to eliminate plasmid DNA. Fragments were eluted using the Zymo DNA Clean and Concentrator-5 Kit.

To generate the barcoded rabies plasmid library for both SAD B19 and CVS-N2c strains, 12 20-μL reactions of gibson assembly were performed for 1 hour at 50C using the HiFi DNA Assembly MasterMix (NEB, E2621), with each reaction containing 184.9 ng of SacII-digested plasmid and a 10-fold molar excess of PCR-amplified insert. The 12 gibson assembly reactions together contain ∼1.3×10^11^ molecules of digested backbone, which is 1000-fold excess to the theoretical complexity of 125 million. The gibson assembly reactions were pooled and purified using Zymo Clean and Concentrator-5 Kit in 12 μL of nuclease-free water pre-heated to 70C. After cooling on ice, the purified gibson assembly reaction was transformed into a total of 300 μL of MegaX DH10B Electrocompetent E. Coli (Thermo, C640003) and grown overnight at 37C in LB-Carbenicillin broth. Some of the transformed bacteria were also plated at 1:1×10^4^, 1:1×10^5^, and 1:1×10^6^ dilution on LB-Agar-Carbenicillin plates, to estimate the total number of colonies to verify sufficient diversity.

The following day, LB-containing bacteria were spun down at 4500×g for 15 min and plasmid DNA was extracted in AE buffer using the Nucleobond 10000 EF Giga Kit (Machery Nagel, 740548) or the ZymoPure II Plasmid GigaPrep Kit (Zymo D4204).

### Generation of barcoded RVdG viral libraries

For library Replicates 1-4 (Figure 1), five 15 cm dishes were coated with 10 μg/mL poly-D-lysine (PDL) diluted in 1× PBS for 1 h at room temperature, and rinsed three times with PBS. Lenti-X HEK293T or HEK-GT^17^ cells that were 70-80% confluent were dissociated using 0.05% Trypsin-EDTA (Thermo, 25200056) and 13 million cells were plated into each PDL-coated 15-cm dish in “FBS-DMEM media”, which was composed of DMEM (Thermo, 41966) containing 10% Heat-inactivated FBS (HI-FBS, Cytiva, SH30071.03HI) and 2% primocin (Invivogen, ant-pm-1). The next day, when the cells had reached ∼90% confluence, each 15 cm dish was transfected with 27.1 μg of SAD B19-N (Addgene #32630), 13.9 μg of SAD B19-P (Addgene #32631), 13.9 μg of SAD B19-L (Addgene #32632), 16.6 μg of CAGGS-T7opt (Addgene #65974), and either 55.1 μg of barcoded pSADdeltaG-dTomato (Libraries 1-2) or pCVS-N2c(deltaG)-MCP-oScarlet-KASH-6XMS2 (Libraries 3-4), using Lipofectamine 3000 (Thermo, L3000075) following the manufacturer’s recommendations. In addition, the Lenti-X HEK293T was co-transfected with 11 μg of SAD B19-G (Addgene #32633). For every microgram of plasmid, 2 μL of P3000 reagent and 2.33 μL of Lipofectamine 3000 was used. The day following transfection, cells were passaged into PDL-coated plates at a 1.28:1 ratio based on surface area (i.e. one 15 cm dish into two 10 cm dishes) using FBS-DMEM media that was additionally supplemented with 1 CEPT cocktail^87^ (50 nM Chroman 1, 5 μM Emricasan, 0.7 μM Trans-Isrib, 1× polyamine supplement). Media was replaced the next day with FBS-DMEM media containing CEPT. Viral supernatant was collected on the fourth, fifth, and sixth day post-transfection to harvest G-pseudotyped virus, with media replacement using FBS-DMEM media containing CEPT. Viral supernatant was collected and filtered through a 0.45 μm syringe filter (Corning 431220). Supernatant was concentrated using PEG precipitation with 1 volume of PEG concentrator for every 3 volumes of supernatant (PEG concentrator: 40% (w/v) PEG 8000 (Promega, V3011) in 1× PBS (Thermo, 10010023) supplemented with 1.2 M NaCl (Fisher, S271-500), and incubated overnight at 4C with gentle agitation. The next day, the precipitated viral particles were spun down by centrifugation at 1600xg for 1 hour at 4C. The supernatant was removed and the viral pellet was resuspended using FBS-DMEM media and 8 μg/mL of polybrene (Millipore Sigma TR-1003-G) at 1:100 of the original media volume.

Per every mL of concentrated G-RVdG virus, 3 million BHK-EnvA-TVA cells^17^ (Library 1), BHK-EnvB cells (Library 2; Salk Institute Viral Core), or BHK-EnvA cells (Libraries 3 and 4; Salk Institute Viral Core) were transduced by spin-infection (1500×g, 1.5 hr, room temperature, using a swinging bucket rotor). Subsequently, cells were plated into 10 cm dishes at a density of 3 million cells/dish in FBS-DMEM media containing 8 μg/mL of polybrene. The following day, the media was replaced with fresh media without polybrene. One or two days later, when the plate reached confluence, cells were passaged once more using 0.25% trypsin-EDTA for 10 minutes at 37C, in order to deactivate any residual G-pseudotyped virus. FBS-DMEM media was added to trypsinized cells at 3-fold volume excess to quench the trypsin, and cells were spun down to remove the trypsin-EDTA. Cells were pooled and passaged at a 1:2.5 dilution (i.e. one 10 cm dish into one 15 cm dish).

Cells were given a media change the next day. Two days later, when plates had reached confluence, viral supernatant was collected every other day for a total of 3-4 collections, depending on the health of the cells. On the day of collection, viral supernatant was filtered through a 0.45 μm syringe filter and concentrated by ultracentrifuge (Beckman Coulter) through a 4 mL sucrose cushion using the SW32Ti rotor (20,300 RPM, 2 hr, 4C). After ultracentrifugation, the viral pellet was resuspended in 50 μL of HBSS while gently rocking overnight at 4C. Virus was aliquoted, frozen at −80C, and titered by infecting TVA800 HEK293T cells (Salk Institute Viral Core) using a serial dilution of virus, and measuring the percentage of dTomato-positive cells two days later by flow cytometry. Titers typically ranged from between 10^7^-10^8^ TU/mL, and lack of G-contamination was confirmed by infecting 100,000 Lenti-X HEK293T cells (that do not express TVA) with 1 μL of undiluted barcoded RVdG virus.

### Testing the effect of cell lines and transfection reagents on RVdG viral rescue efficiency

To compare rescue efficiency in different packaging cell lines, Lenti-X HEK293T cells, BHK-21, and Neuro2A cells were passaged into PDL-coated 24-well plates as described above at a density of 1.75 x 10^5^ cells per well (HEK293T, Neuro2A) or 1.1 x 10^5^ cells per well (BHK-21). The next day, when the cells had reached ∼90% confluence, each well was transfected with 760 ng of barcoded pSADdeltaG-dTomato, 374 ng of SAD B19-N, 192 ng of SAD B19-P, 192 ng of SAD B19-L, and 229 ng of CAGGS-T7opt using Lipofectamine 3000 following manufacturer recommendations. For every μg of DNA, 2 μL of P3000 reagent and 2.33 μL of Lipofectamine 3000 were used. The next day, the media was changed with fresh FBS-DMEM media. Two days later, cells were dissociated using 0.05% Trypsin-EDTA, quenched with three volumes of FBS-DMEM media for every one volume of Trypsin-EDTA, and pelleted by centrifugation at 300×g for 3 minutes. Cells were resuspended using 1× PBS containing 0.1% bovine serum albumin (BSA) and filtered into a flow cytometry tube through a 35 μm nylon mesh cap. Flow cytometry was performed while recording from 50,000 events to quantify the percentage of cells that had successfully recovered RVdG viral particles based on dTomato expression.

To compare rescue efficiency in HEK293T comparing different transfection reagents, Lenti-X HEK293T cells were passaged into PDL-coated 24-well plates as described above, and transfected using Lipofectamine 3000 as described above, Lipofectamine 2000 following published recommendations^14^ (1.17 μL per 1 μg of DNA), XFect following published recommendations^4^ (0.3 μL per 1 μg of DNA), FuGENE HD (3 μL per 1 μg of DNA), PEI MAX (3 μL per 1 μg of DNA), XtremeGene HP (3 μL per 1 μg of DNA), and Mirus TransIT-293 (3 μL per 1 μg of DNA). Media was changed the next day and flow cytometry was performed three days after transfection.

### Production of barcoded RVdG for comparison of transfection reagents, plate size, and cell lines

To compare the effect of packaging RVdG using Lipofectamine 2000 in HEK293T cells, Lipofectamine 3000 in HEK293T cells, and Lipofectamine 3000 in HEK-GT cells on RVdG viral library complexity, eight PDL-coated 15-cm dishes were plated with Lenti-X HEK293T cells and four PDL-coated 15-cm dishes were plated with HEK-GT cells as described above. Each dish functioned as a technical replicate for this experiment. Each 15 cm dish was transfected with 55.1 μg of barcoded CVS-N2c MCP-oScarlet-KASH, 27.1 of SAD B19-N, 13.9 μg of SAD B19-P, 13.9 μg of SAD B19-L, and 16.6 μg of CAGGS-T7opt using Lipofectamine 3000 or Lipofectamine 2000 as described above. Transfection of HEK293T, but not HEK-GT cells additionally included 11 μg of SAD B19-G. The next day, the contents of each 15 cm dish were passaged using 0.05% Trypsin-EDTA into two 10-cm dishes, while keeping cells from each 15 dish separate. Media was changed the next day, and supernatant was collected to harvest G-pseudotyped RVdG when approximately half of the cells were oScarlet-positive (day 5 post-transfection for Lipofectamine 3000 in HEK293T and HEK-GT cells, day 8 post-transfection for Lipofectamine 2000). The supernatant from the 10-cm dishes that originated from the same 15-cm dish were pooled and concentrated using PEG precipitation, and viral genome RNA was isolated as described in the next section.

To compare the effect of packaging RVdG in 10-cm dishes and 384 well plates, two PDL-coated 15-cm dishes containing LentiX HEK293T cells were transfected with packaging plasmids as described above. The next day, cells from one 15-cm dish were passaged into two PDL-coated 10-cm dishes, and cells from the other 15-cm dish were passaged into five PDL-coated 384 well plates. Media was changed the next day and supernatant was harvested four days post-transfection for concentration and viral genome RNA isolation as described in the next section.

### Generation of sequencing library for plasmid and viral barcode diversity quantification

To quantify plasmid barcode library diversity, 50 ng of plasmid DNA was tagmented using seqWell’s Tagify UMI reagent, in order to tag each barcode-containing plasmid DNA fragment with a UMI and partial Illumina Read1 adapter. Tagmented fragments were purified using a 1× left-sided size selection using SPRIselect beads according to the manufacturer’s protocol. Sample indexing PCR was subsequently performed using custom sample index primers that bind to the TruSeq P5 PCR handle generated during Tn5 tagmentation (Primer 3, Table S2) and to the BFP primer site sequence immediately proximal to the barcode sequence (Primers 8-15, Table S2). The library was sequenced using a target of 100 million reads.

To quantify viral barcode library diversity, barcoded RVdG from a single 15 cm dish was concentrated by ultracentrifugation or PEG precipitation as described above and RNA was isolated by phenol-chloroform extraction, the procedure for which is described as follows. The viral pellet was resuspended in 1 mL of Trizol (Invitrogen 15596026), and incubated at room temperature for five minutes to permit dissociation of rabies proteins from viral RNA. Subsequently, 200 μL of chloroform (Fisher 60047878) was added and vortexed for 30 seconds, followed by spin at 12000×g for 15 min at 4C. 600 μL of the upper aqueous phase was collected and RNA was precipitated by mixing with 500 μL of isopropanol and 1 μL of Glycoblue (Invitrogen AM9515) and incubating for 10 minutes. Precipitated RNA was pelleted by centrifugation at 12000×g for 10 min at 4C, washed twice with 80% ethanol, air dried for 2 min, and resuspended in 10 μL of nuclease-free water. Reverse transcription was performed on 1-5 μg of RNA using SuperScript IV (ThermoFisher 18090010) and 1 μL of 50 mM custom RT primer (Primer 4, Table S2) targeting a sequence distal to the barcode relative to the negative stranded viral genome, in order to generate cDNA containing barcodes from viral genome RNA but not from contaminant mRNA. RT primer also contained an overhanging sequence carrying an 8 bp randomer, which functions as a UMI to correct for PCR amplification bias after sequencing, and a binding site for Illumina sample index primer. The resulting cDNA (5 uL, or 25% of the RT reaction) was amplified using 10-12 cycles of PCR with KAPA HotStart PCR ReadyMix (Roche KK2601) and custom sample index primers (P7 primers: Primers 5-7; P5 primers: Primers 16-23; Table S2), one of which is specific to an overhang introduced by the RT primer to facilitate viral genome-specific amplification. Each library was sequenced using at least 10 million reads.

### Organotypic slice culture and viral transduction of human brain

To generate organotypic slice cultures, cortical tissue was identified by visual inspection, embedded in 3% low melting point agarose (Fisher, # BP165–25), and sliced into 300 μm sections perpendicular to the ventricle using a Leica VT1200S vibrating blade microtome. Slicing was performed in oxygenated artificial cerebrospinal fluid (ACSF) containing 125 mM NaCl, 2.5 mM KCl, 1 mM MgCl_2_, 1 mM CaCl_2_, 1.25 mM NaH_2_PO_4_. Slices were cultured on Millicell Standing Cell Culture Inserts (PICM03050) in a serum-free media composed of DMEM-F12 with Glutamax (Gibco 10565018) and Neurobasal media (Gibco 21103049) mixed at a 1:1 (v/v) ratio and supplemented with 2% B27 (Gibco 17504044), 1% N2 (Gibco 17502048), 200 μM ascorbic acid-2-phosphate (Wako 50-990-141), and 2% primocin. One hour after plating, slices were transduced with a VSV-G pseudotyped lentiviral helper virus (pSico-hSyn1::GFP-T2A-TVA-E2A-oG) described above by pipetting 2 μL of 10^6^ TU/mL over the top of the slice. Half of the media was exchanged every other day subsequently. Slices were transduced with barcoded RVdG-dTomato by pipetting 3 uL of 10^7^ TU/mL virus over the top of the slice three days after the initial helper virus infection. Five days after transduction with RVdG-dTomato, slices were dissociated into a single cell or single nuclei suspension following previously published protocols^88,89^. For single cell dissociation, 100 units of papain in 1 mM L-cysteine and 0.5 mM EDTA were reconstituted in 5 mL of EBSS containing 5% Trehalose and reactivated for 10 minutes. 500 units of DNAse I was added to 5 mL of reactivated papain solution and slices were incubated in 1 mL of papain/slice for 30 min at 37C. Papain was subsequently inactivated using EBSS containing 58.3% ovomucoid inhibitor, 5% DNAse I, and 5% Trehalose (EBSS-ovomucoid), using a 1:1 (v/v) papain:EBSS-ovomucoid inhibitor ratio. Slices were spun down at 100×g for 10 min at 4C, resuspended in 1 mL of EBSS-ovomucoid, and gently triturated 30 times using a fire-polished glass pipette, or until the tissue has been fully dissociated into a single cell suspension. 4 mL of PBS containing 1% BSA, 5% Trehalose, and 5% DNAse I was added per 1 mL of EBSS-ovomucoid, mixed gently, and spun down at 100×g for 10 min at 4C. Cells were resuspended in ice-cold PBS containing 1% BSA and filtered through a 35 μm strainer mesh in preparation for fluorescence-activated cell sorting (FACS).

For single nuclei dissociation, slices were placed in a pre-chilled 1 mL dounce homogenizer (Wheaton EW-04468-00) and slowly dounced ten times with the loose pestle A and ten times with the tight dounce pestle B while on ice using 1 mL EZ lysis buffer (Sigma NUC-101) supplemented with 5 μL of RNAse inhibitor (Takara 2313B; 40 U/uL). The contents of the dounce were moved to a pre-chilled 15 ml conical tube, incubated for 5 min on ice, and spun down in a swinging bucket centrifuge at 4C, 500×g, for 5 min.

Supernatant was gently removed, and nuclei were gently resuspended in ice-cold 1 mL EZ lysis buffer supplemented with 5 μL of RNAse inhibitor using 3-5 triturations using a 1 mL pipet and normal-bore tips.

Nuclei were incubated for 5 min, spun down in a swinging bucket centrifuge at 4C, 500×g, for 5 min, and resuspended in ice-cold PBS containing 1% BSA and filtered through a 35 μm strainer mesh in preparation for fluorescence-activated nuclei sorting (FANS).

FACS/FANS was performed to isolate dTomato+ cells or oScarlet+ nuclei using a 100 μm or 85 μm nozzle, respectively, on a BD FACSAria Fusion. Cells were collected in a PCR tube nested within a 600 uL tube nested within a 1.5 mL Eppendorf tube^90^ containing 50 μL of collection buffer consisting of PBS-1% BSA and 200 units/mL RNAse inhibitor. The PCR tube had previously been precoated with 1% BSA for one hour, to prevent cells from sticking to the side of the tube during FACS/FANS. One final centrifugation was performed to pellet the cells/nuclei (cells: 300×g for 5 min, nuclei: 500×g for 5 min), and the supernatant was carefully removed to leave behind approximately 4 μL of collection buffer.

### Generation of transcriptome and barcode libraries

scRNA-seq and snRNA-seq was performed on collected cells/nuclei using the PIPseq v4.0 T2 3’ Single Cell RNA kit, adhering to the manufacturer’s protocol for isolating single-cell and single-nucleus transcriptomes. Transcriptomes were sequenced with a target of 35,000 reads/cell.

For generating barcode libraries, PCR was performed on 25% of the cDNA amplification library using custom sample index primers that target the BFP primer site immediately proximal to the barcodes (Primers 8-15; Table S2) and the partial Illumina Read1 sequence introduced by the 10x RT primer (Primers 16-23; Table S2). 10-15 rounds of PCR was performed using the following program: 1) 98C, 30s, 2) 98C, 10 sec, 3) 62C, 20 sec, 4) 72C, 10 sec, 5) Repeat steps 2–4 9-14×, 6) 72C, 2 min, 7) 4C, hold, as described previously^11^. Barcode library was purified by 0.6-1× dual-sided size selection using SPRIselect beads (Beckman Coulter B23317) and sequenced with a target of 5,000-10,000 reads/cell using the following sequencing parameters: Read 1: ≥ 54 reads, Index 1: 8 reads, Index 2: 8 reads, Read 2: 150 reads.

### Whole-cell patch-clamp recordings

Electrophysiological experiments were conducted on cultured human prenatal subplate neurons at DIV 6-8 (i.e. 3-5 days after transduction with RVdG-dTomato). Cultured brain slices were placed in a recording chamber mounted on a BX51WI microscope (Olympus) and were perfused with artificial cerebrospinal fluid (ACSF) containing (in mM): 125 NaCl, 2.5 KCl, 1.25 Na H_2_PO_4_, 25 NaHCO_3_, 11.1 Glucose, 1 MgCl_2_, 2 CaCl_2_, pH 7.4, 305-310 mOsm, equilibrated with 95% O_2_, 5% CO_2_ at 34 ± 1°C. Fluorescence imaging with a 40× water immersion objective lens (LUMPLFLN40XW, NA 0.8) was used to identify starter and non-starter cells (GFP and dTomato/oScarlet fluorescence together vs. dTomato/oScarlet fluorescence alone, respectively). Whole-cell patch-clamp recordings were performed on fluorescently labeled cells using 6–8 MΩ resistance glass microelectrodes (BF150-86-10; Sutter Instrument) containing an internal solution comprised of (in mM): 120 K gluconate, 4 KCl, 10 HEPES, 4 MgATP, 0.3 Na_3_GTP, 10 sodium phosphocreatine, 13.4 biocytin, pH 7.25 with 0.5M KOH, ∼320 mOsm. In a subset of experiments, a slightly modified internal solution was used consisting of (in mM):111 K gluconate, 4 KCl, 10 HEPES, 0.2 EGTA, 4 MgATP, 0.3 Na_3_GTP, 5 sodium phosphocreatine, 13.4 biocytin, pH 7.25 with 0.5M KOH with addition of 1 U/μl recombinant RNase inhibitor (Roche) immediately prior to experiment and final osmolarity of 315–320 mOsm. All signals were obtained using Multiclamp 700B amplifier and Digidata 1550B (Molecular Devices). Specifically, voltage-clamp recordings were sampled at 20 kHz, low-pass filtered at 2kHz, and current-clamp recordings were sampled at 50 kHz, low-pass filtered at 10 kHz. Easy Electrophysiology software (Easy Electrophysiology Ltd.) was used to analyze recorded traces that met the following quality control criteria: (1) >1 GΩ seal before whole-cell configuration; (2) series resistance <25 MΩ; and (3) <20% series resistance changes between before, during and after the recording. In current clamp configuration, 1-s long current steps were injected ranging from −20 pA to 60 pA in 2 pA increments to elicit action potentials. The resting membrane potential was measured as the mean membrane potential during 100 ms before the current stimulation. The input resistance was calculated as the voltage difference between the steady state and the resting membrane potential divided by the injected current during the hyperpolarizing steps. The cell membrane capacitance was measured as the time constant of a monoexponential fit to the first local minimum of the hyperpolarizing membrane potential during stimulation divided by the membrane resistance. Action potential was identified initially by locating the action potential threshold, which was defined as the potential at which the first derivative of the membrane potential (dV/dt) exceeded 20 mV/ms. Only those peaks with the following criteria were detected as an action potential: > 20 mV amplitude, > 20 mV/ms upstroke, > −0.1 mV/ms downstroke, < 5 ms width.

## QUANTIFICATION AND STATISTICAL ANALYSIS

Statistical analyses were performed in RStudio and Jupyter. Information regarding statistics including the statistical test, sample sizes, and p-values for each experiment are provided in the figure legends and the Results section. A p-value of less than 0.05 was considered statistically significant, with corrections applied for multiple comparisons when necessary.

### Viral barcode pre-processing

Output fastqs from sequenced rabies barcode libraries were filtered and processed using the custom script barcode_processing.sh. The bbduk^91^ (v39.01) package (BBMap) was used to discard any reads with average quality (maq) below 20. Reads were then trimmed down to individual RVdG barcodes, UMIs, and cell barcodes (for transcriptome-matched datasets). RVdG barcodes were aligned to a custom reference white list (Table S1) with bowtie2, allowing for one base pair mismatch per read. After sorting output files with samtools, the RVdG barcodes, UMIs, and cell barcodes were merged on read IDs. The command line package UMI-tools^92^ (v1.1.2) was then used to quantify the number of UMIs per cell barcode and RVdG barcode combination. A UMI collapsing threshold was set at 1 to collapse any UMIs with only one base pair difference into a single UMI count.

### Viral barcode diversity analysis

When available, UMI counts per RVdG barcode were retrieved from previously published studies^3–5^. For those studies that were not already in the output format of UMI-tools, counts per barcode were generated using UMI-tools from published barcode lists. Barcode library diversity was compared using the percentage of the UMIs covered by each unique rabies barcode per dataset plotted with Matplotlib^93^ (v3.9.1).

### Barcode collision simulations

To evaluate the risk of barcode collision (or the chance of a cell receiving the same barcode as a previously infected cell), we set up a simulation in which 10-100,000 cells (for viral libraries) or 10-10,000,000 cells (for plasmid libraries) are assigned a single barcode by bootstrapping from the original dataset with replacement (Figures 1E, S1K, S1M, and S1O). Each simulation was repeated 50 times using numpy.random.choice in the Numpy^94^ package, and duplicate barcodes were counted with the pandas^95^ package’s pandas.dataframe.duplicated(keep=first) command to ensure that barcode collision was counted starting from the first duplicate barcode infection. Barcode collision was then plotted as a function of the number of cells per simulation. Line plots showing the collision metric value at each of the starter cell counts were drawn using the Seaborn^96^ (v0.12.2) package in Python with error bars extended to two standard error widths based on the results from the 50 iterations for each simulation.

### Barcode sampling simulations

To examine the total number of unique barcode as a function of the number of transcripts sequenced, we set up a simulate in which 10-1,000,000 barcodes were pulled from barcode diversity datasets at random using using numpy.random.choice in the Numpy^94^ with replacement (Figure 1F). These simulations were repeated 100 times, and error was calculated as standard deviation. Line plots showing the total number of unique barcodes pulled versus the total number of all barcodes pulled were plotted using Matplotlib^93^ (v3.9.1) with error bars extending to two standard error widths based on the results from the 100 iterations for each simulation.

### Shannon entropy calculations

To estimate the Shannon entropy for the libraries presented and previous published ones, the function shannon_entropy_calculator() was created. Iterating through a list of barcode libraries, it computes the observed barcode probability distributions by dividing the UMI count of a barcode by the sum of UMI counts in the library. Uniform barcode probability distributions are calculated by specifying each barcode probability as 1 divided by the sum of UMI counts in the library. The probability distributions are inputted into the SciPy entropy() function to calculate the observed and uniform Shannon entropies (in bits) for each library.

### Transcriptomic processing and analysis

Output fastqs from PIPseq datasets were first aligned using the PIPseeker^3^ (v3.3) prebundled hg38 genome to generate mitochondrial gene counts and complete cell barcodes. The argument –retain-barcoded-fastqs was used to retain original fastqs with modified cell barcodes. These fastqs were then concatenated per dataset and realigned with STAR (v2.7.11b) to the hg38 reference genome with NCBIRefSeq annotations to generate transcriptomic count matrices. Filtered matrices with the lowest sensitivity were processed using Seurat (v4.3)^97^. Briefly, single-cell datasets were filtered for cells with >1000 features, >1250 UMI counts, and <5% mitochondrial genes. Single-nuclei datasets were filtered for nuclei with >300 features, >300 UMI counts, and <5% mitochondrial genes. To identify cell types, all datasets were merged and normalized using SCTransform(). The top 2000 variable features were selected, and mitochondrial genes were regressed out from further analysis. PCA was calculated using the top 50 principal components of the SCT assay. The package Harmony^22^ (v1.2.0) was then used to integrate all datasets to account for batch effects and differences between dissociation methods. RunUMAP(), FindNeighbors(), and FindClusters() were used to cluster the resultant object, accounting for dimensions 1:40 with cluster resolution set to 1.0. Marker genes across clusters were identified using the package presto^98^ (v1.0.0) and Seurat’s FindAllMarkers() function. Cell type labels were identified based on expression of key marker genes. These cell type labels were retained for all downstream analyses.

### Gene set enrichment analysis

To identify differentially expressed genes between the uninfected, SAD B19-, and CVS-infected datasets dissociated as single-cells, these datasets were merged and then processed using the same parameters described above, excluding integration with Harmony. Differentially expressed genes between uninfected, SAD B19-, and CVS-infected datasets and each cell type were identified using Seurat’s FindMarkers() function. Top differentially expressed genes were visualized using the EnhancedVolcano^99^ (v1.20.0) package. Gene set enrichment analysis was conducted using the gseGO() function in clusterProfiler^100,101^ (v4.10.1) and restricted to the Gene Ontology’s Biological Processes database. FDR adjusted p-values were reported, and the top ten pathways with an adjusted p-value <0.05 were visualized using the dotplot() function. Cell type-specific enrichment of certain terms was confirmed by calculating term enrichment per cell using the AddModuleScore_UCell() function in the UCell^102^ package (v2.6.2), then mapping enrichment scores using Seurat’s FeaturePlot() function.

### Nuclear and cellular viral barcode metric comparisons

Medians and distributions of UMI counts for every unique rabies barcode found in every unique cell/nucleus, as well as the total number of rabies barcode UMIs per every unique cell/nucleus were plotted using Matplotlib (v3.9). The relationship between total rabies barcode UMIs and total unique rabies barcode per cell/nucleus and lines of best fit were plotted with seaborn (v0.13.2). R^2^ was calculated by squaring the Spearman’s Correlation coefficients calculated using SciPy’s (v1.11.1) spearmanr() function.

### Empty droplet analysis

Cell/nucleus barcodes representing empty droplets were identified from Pipseeker’s generated barcode read info table as any cell barcode with ≤10 transcripts mapping to the hg38 reference genome. All rabies barcodes with cell/nucleus barcodes matching those corresponding to empty droplets were considered for empty droplet analysis. The distribution and central tendencies of UMIs per rabies barcode per cell/nucleus or empty droplet, total rabies barcode transcripts, and unique rabies barcode identified across either real cells/nuclei or empty droplets were calculated and visualized using SciPy’s^103^ (v1.11.1) ranksums() function with Bonferroni correction and Matplotlib packages. Fractions of barcodes in empty droplets, cells/nuclei, or both were visualized using ggplot2 (v3.5.1).

### Preparing barcode data for connectivity analysis

Barcode data was prepared for connectivity analysis using the function process_barcodes_df() written in Python (Table S4). This filtered for cells that passed QC, and annotated each RVdG barcode-cell/nucleus barcode (CBC) based on the CBC’s cell type and number of helper UMIs. Barcode and helper UMI thresholds of ≥3 were used to filter for barcodes and assign starter status. Additional annotations were made for barcodes that were found in at least 2 cells; whether the barcode was found in starter cells, non-starters, or both; and whether the barcode was only found in one starter cell (termed “single_starter_barcodes”).

### Identification of single-starter networks

To systematically infer single-starter networks involving a single starter cell and all of the non-starter cells that share at least one one barcode with the starter, we filtered the dataset generated using the function process_barcodes_df() for single starter barcodes that were found in at least one nonstarter cell. All of such barcodes associated with a cell were grouped into a set for each cell. Iterating through each starter cell’s barcode set and all of the non-starter cell’s barcode sets, non-starters that shared at least one barcode with a given starter cell (as determined by the intersection of their sets being >0) were matched with that starter cell’s network. By running this algorithm for all starter cells that have at least one single-starter barcode, two connectivity datasets are generated: 1) a dataset with each row corresponding to a starter and aggregated information on its connections 2) a dataset outlining, for each connection, the CBCs and cell types of the starter and non-starter cells.

### Network visualization and analysis

To visualize the outputs from connectivity inference, graph creation functions (graph_draw() and graph_plot()) were created to convert the datasets into directed graphs using the NetworkX package^104^ in Python. Graphs were drawn using the following customizations: Neato layout using Graphviz^105^ for node positioning, node size scaled by degree, nodes colored by cell type, edges with uniform thickness, and edges

### Analysis of cell type–specific connectivity enrichment

To identify biases in observed cell type connectivity and how they differ from networks predicted by chance, observed and null connectivity matrices were generated. By aggregating data from single-starter networks by cell type, an observed adjacency matrix was calculated with rows and columns corresponding to the non-starter and starter cell types, respectively. To generate the observed connectivity matrix, we divided the number of connections observed between each pair of cell types (e.g. non-starter GABAergic to starter L2/3 ExNs) by the total number of connections observed. The null connectivity matrix was determined by the relative abundance of cell types among our starter cell population (starter cell probabilities) and cell types in our slice overall based on the uninfected population (non-starter cell probabilities). Starter cells were defined as cells from all of the combined SAD B19 (n=7 datasets) and CVS-N2c (n=4 datasets) datasets described above with ≥3 UMIs of helper virus transcripts detected. In addition, we generated a dataset in which the helper virus expressed GFP and TVA but not G, and included all of the dTomato+ cells from this experiment as starter cells for this analysis (Table S3). Non-starter probabilities were derived from the population of uninfected cells. The null connectivity matrix was then generated by multiplying the proportion of each cell type among non-starter cells in a pairwise manner by the proportion of each cell type among starter cells.

Enrichment scores were computed by dividing each observed probability by the corresponding null probability and log10 scaling the ratios to center them around 1, with negative values indicating the observed connection was less likely than chance and positive values indicating the connection was more likely than chance. To assess for statistically significant differences between the observed and null probabilities, the null distributions for each connection were modeled using binomial distributions (n = total number of observed connections, p = null probability of connection between two cell types) and two-sided binomial tests using the SciPy package in Python with Bonferroni connections were performed. P-values less than .05 were considered significant.

## SUPPLEMENTAL INFORMATION

Document S1. Figures S1-S8

Table S1. Barcode whitelist. Data from a recent publication^11^, related to Figures 1 and S1 Table S2. Primers used for sequencing library generation, related to Figures 1, 2, 6, 7, and S1 Table S3. sc/snRNA-seq metadata, related to Figures 4, 5, 6, 7, 8 and S4, S5, S7, and S8 Table S4. Cell type-matched RVdG barcodes for connectivity analysis, related to Figures 8, S7, and S8

