## Supplemental file for "High-Complexity Barcoded Rabies Virus for Scalable Circuit Mapping Using Single-Cell and Single-Nucleus Sequencing"

**SUPPLEMENTAL INFORMATION**

Supplementary Figures S1 – S8

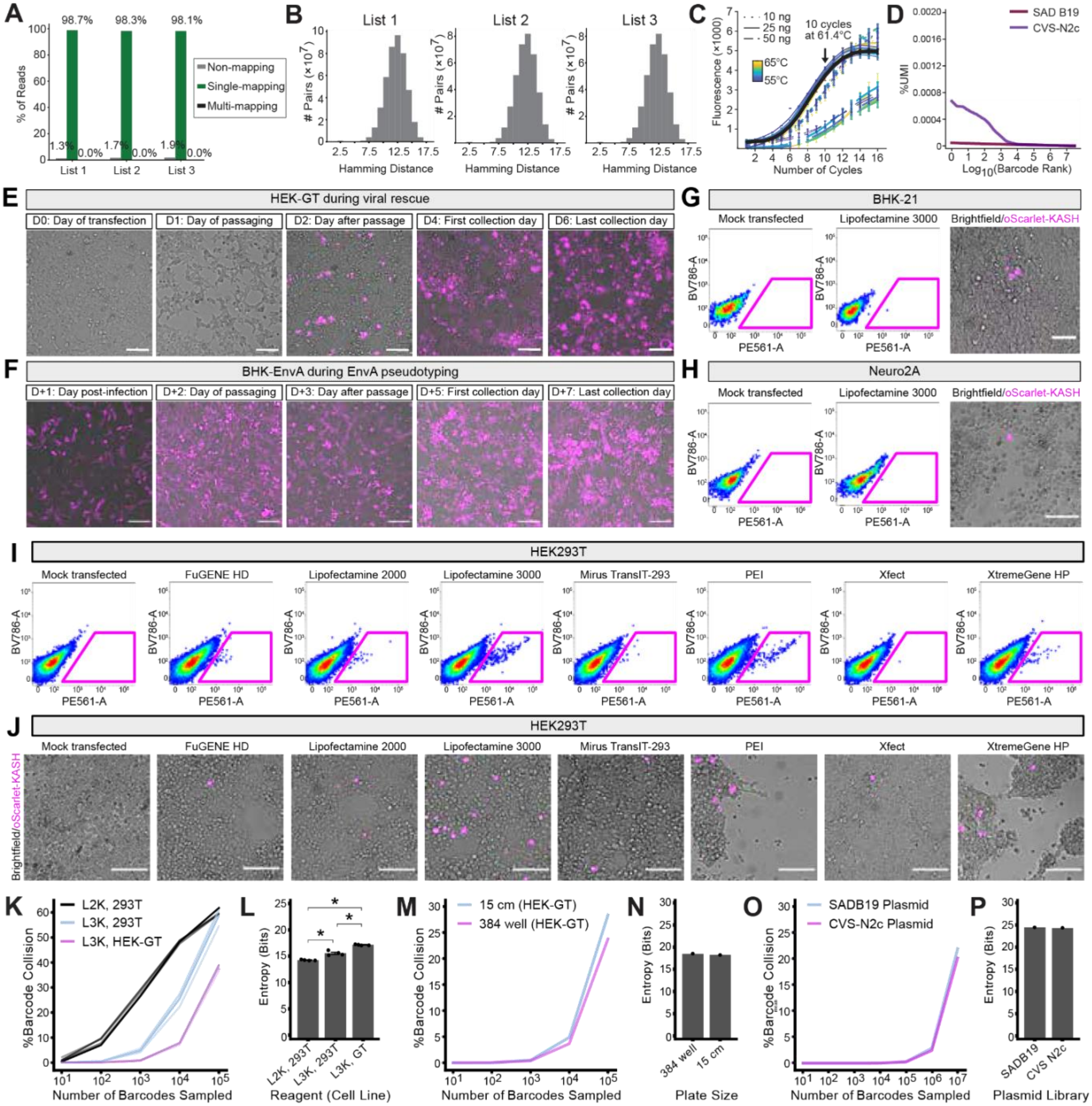


**Supplemental Figure 1. Related to Figures 1 and 2. Quality control validation of barcoded RVdG plasmid and viral libraries and optimizations to improve barcode diversity. A.** Fraction of barcode reads mapping to a single barcode, multiple barcodes, or not mapping. **B.** Actual hamming distance between sequenced barcodes in the packaged viral library. **C.** KAPA qPCR optimization of the number of cycles sufficient to amplify the barcode fragment for cloning into the RVdG plasmid. **D.** Barcode diversity in SAD B19 and CVS-N2c viral genome plasmid libraries. **E.** Flow cytometry gating to determine the percent of cells that successfully rescued RVdG viral particles three days after transfecting with packaging plasmids in the absence of rabies glycoprotein G. **F.** Visualization of fluorescent cells three days after transfecting with packaging plasmids in the absence of rabies G. **G-H.** Flow cytometry gating and visualization of fluorescent cells on day 3 of packaging in the absence of rabies G in BHK-21 **(G)** or Neuro2A cells **(H)**. **I-J.** Representative images during the two stages of rabies packaging: transfection and viral propagation in HEK-GT cells **(I)** and EnvA-pseudotyping in BHK-EnvA cells **(J)**. **K.** Percentage of simulated barcode collision events, or at least two cells receiving the same barcode by chance, as a function of the number of infections simulated comparing transfection of HEK293T cells using Lipofectamine 2000 (Lipo2000, 293T) or Lipofectamine 3000 (Lipo3000, 293T) and HEK-GT cells using Lipofectamine 3000 (Lipo3000, GT). Error bars are 95% confidence intervals for 50 iterations. **L.** Estimate of Shannon entropy for comparison in **(K)**. Pairwise Student's t-test with Bonferroni-Holm correction was performed (n = 4 x 15 cm dishes per experimental condition), *$p<0.05$. **M-N.** Percentage of barcode collision events as a function of number of barcodes sampled **(M)** and Shannon entropy **(N)** comparing packaging in HEK-GT using different plate sizes at equivalent surface area. **O-P.** Percentage of barcode collision events as a function of number of barcodes sampled **(O)** and Shannon entropy **(P)** comparing barcoded SAD B19 and CVS-N2c RVdG plasmid libraries.

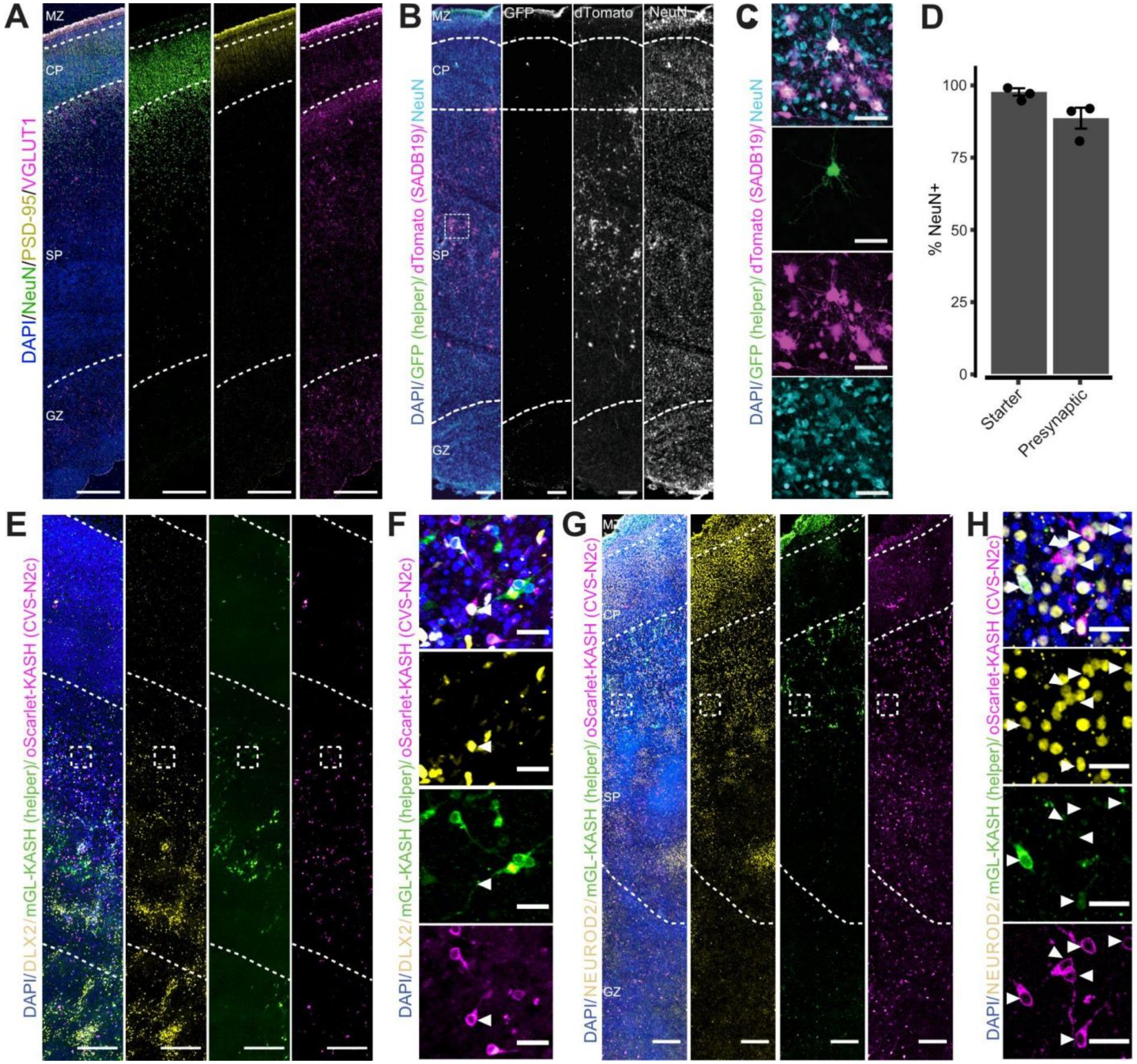


**Supplemental Figure 2. Related to Figure 3. Monosynaptic rabies tracing using SAD B19 and CVS-N2c strain RVdG in human prenatal organotypic slice cultures. A.** Full thickness tile scan of GW22 cortex stained for pan-neuronal marker NeuN, presynaptic glutamatergic marker VGLUT1, and postsynaptic marker PSD95. Scale bars, 250 μm. **B-D.** Tile scan **(B)**, high magnification image **(C)**, and quantification **(D)** of slice stained for GFP (cells infected with helper virus), RFP (cells infected with SAD B19 RVdG), and NeuN (n=3 slices from 3 individuals). **E-H.** Images of slices infected with CVS-N2c strain RVdG and stained for GFP (cells infected with helper virus), RFP (cells infected with oScarlet-KASH), and DLX2 **(E-F)** or NeuroD2 **(G-H)**. Scale bar for tile scan, 250 μm; for high magnification: 25 μm.

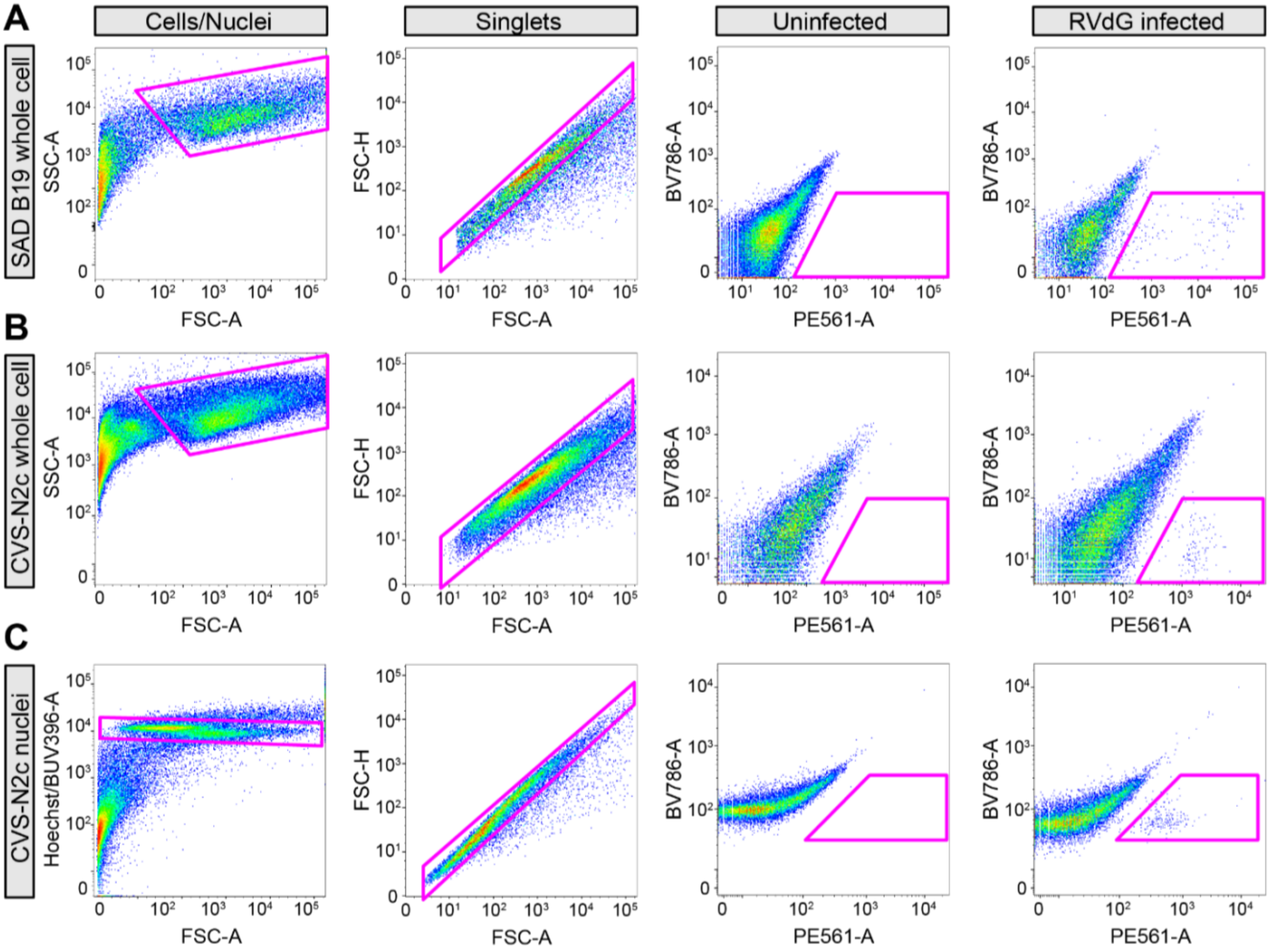


**Supplemental Figure 3. Related to Figures 4-6. Gating scheme for fluorescence activated cell/nuclei sorts (FACS/FANS) for single cell/nuclei sequencing. A.** FACS gating scheme after whole cell dissociations of tissue infected with SAD B19. **B.** FACS gating scheme after whole cell dissociations of tissue infected with CVS-N2c. **C.** FACS gating scheme after nuclei isolations of tissue infected with CVS-N2c.

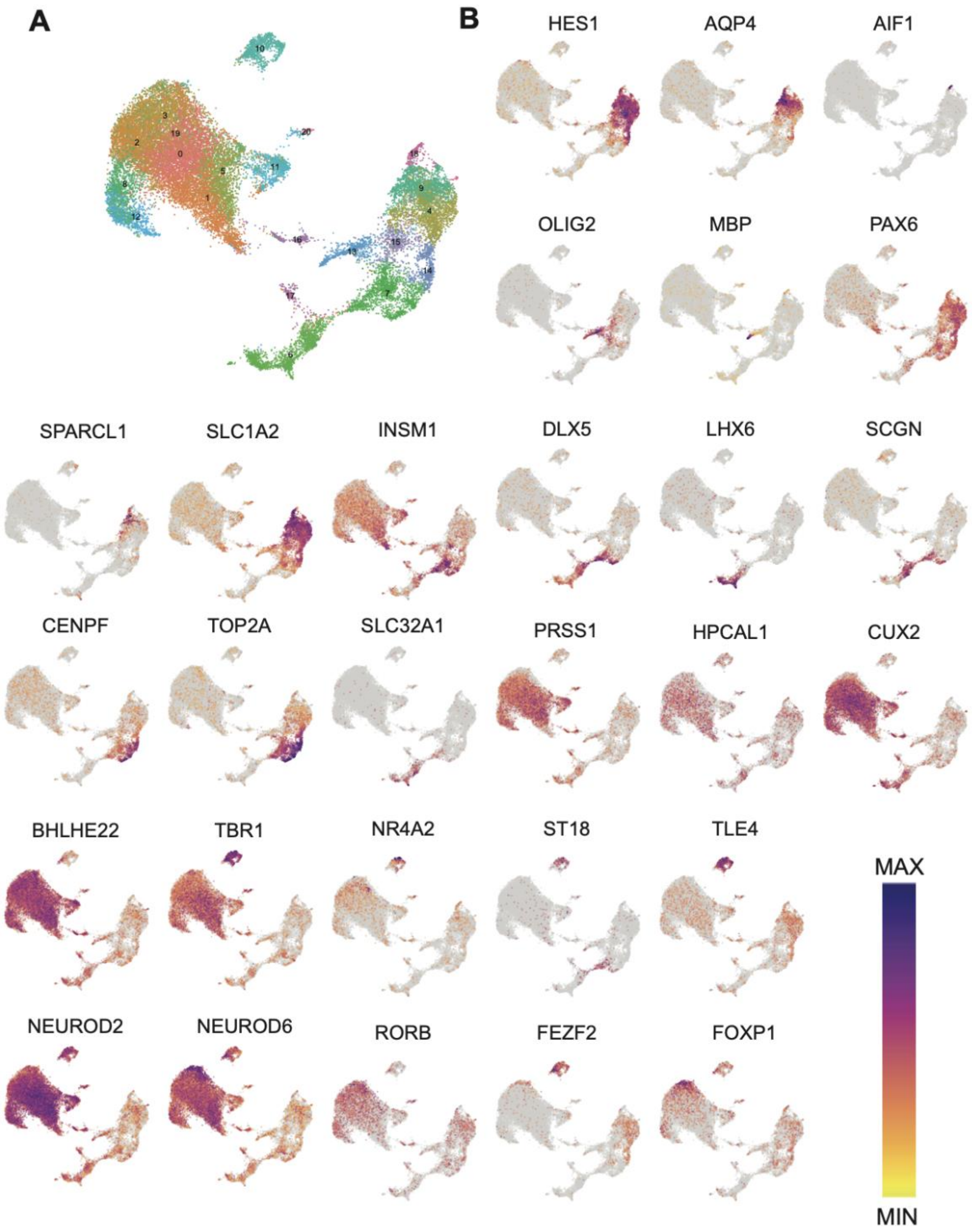


**Supplemental Figure 4. Related to Figure 4. Cell type identification of pooled RVdG datasets. A.** UMAP plots with original clusters. **B.** Feature plots of key marker genes used to annotate cell types. Color is scaled relative to expression level of that gene across all cells in pooled datasets.

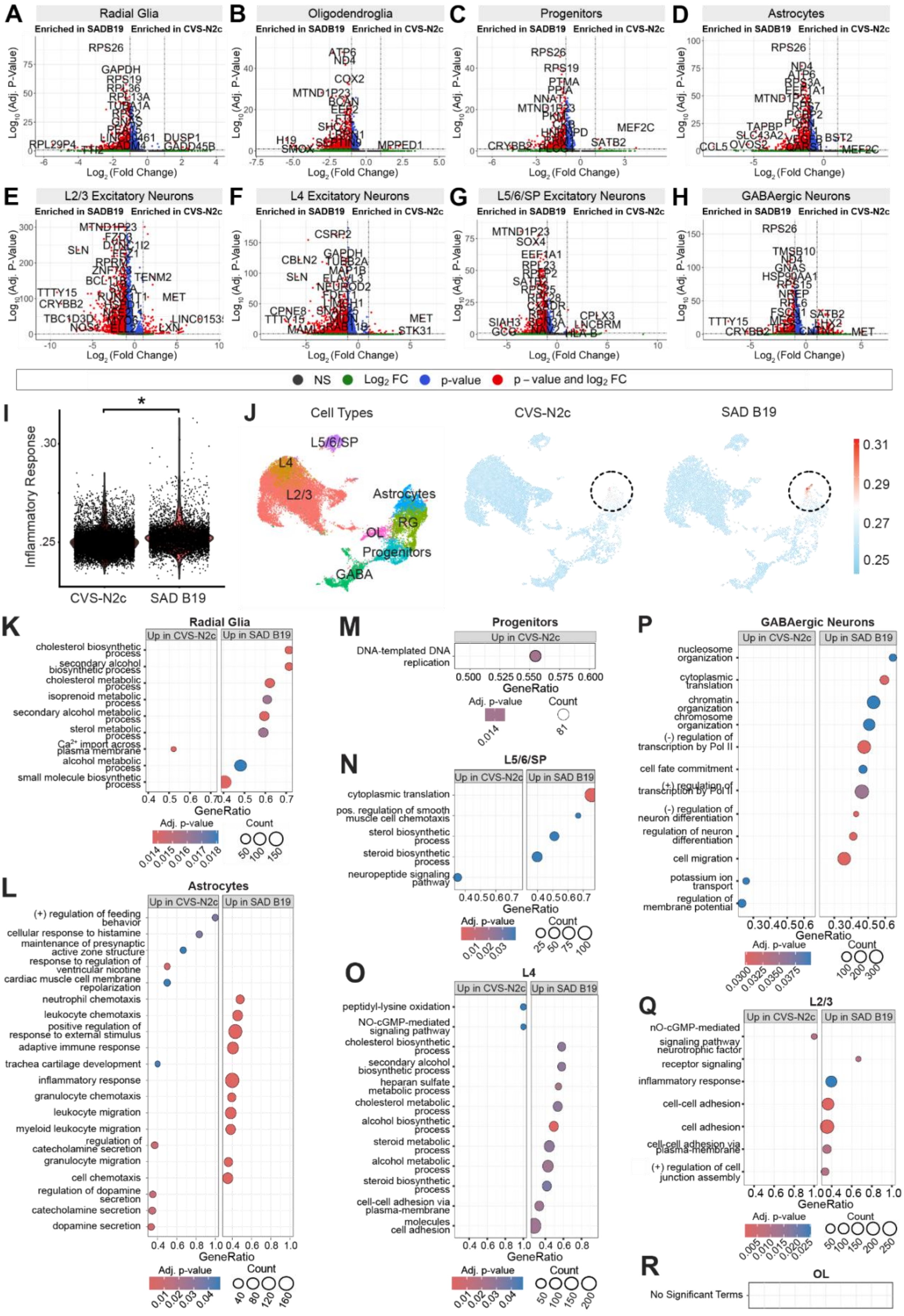
A
Radial Glia
B
Oligodendroglia
C
Progenitors
D
Astrocytes
Enriched in SADB19
Enriched in CVS-N2c
Log10 (Adj. P-Value)
Log2 (Fold Change)
E
L2/3 Excitatory Neurons
F
L4 Excitatory Neurons
G
L5/6/SP Excitatory Neurons
H
GABAergic Neurons
NS
Log2 FC
p-value
p – value and log2 FC
I
Inflammatory Response
CVS-N2c
SAD B19
J
Cell Types
L5/6/SP
L4
L2/3
Astrocytes
RG
OL
Progenitors
GABA
CVS-N2c
SAD B19
K
Radial Glia
Up in CVS-N2c
Up in SAD B19
cholesterol biosynthetic process
secondary alcohol biosynthetic process
cholesterol metabolic process
isoprenoid metabolic process
secondary alcohol metabolic process
sterol metabolic process
Ca2+ import across plasma membrane
alcohol metabolic process
small molecule biosynthetic process
GeneRatio
Adj. p-value
Count
M
Progenitors
Up in CVS-N2c
DNA-templated DNA replication
GeneRatio
Adj. p-value
Count
P
GABAergic Neurons
Up in CVS-N2c
Up in SAD B19
nucleosome organization
cytoplasmic translation
chromatin organization
chromosome organization
(-) regulation of transcription by Pol II
cell fate commitment
(+) regulation of transcription by Pol II
(-) regulation of neuron differentiation
regulation of neuron differentiation
cell migration
potassium ion transport
regulation of membrane potential
GeneRatio
Adj. p-value
Count
N
L5/6/SP
Up in CVS-N2c
Up in SAD B19
cytoplasmic translation
pos. regulation of smooth muscle cell chemotaxis
sterol biosynthetic process
steroid biosynthetic process
neuropeptide signaling pathway
GeneRatio
Adj. p-value
Count
L
Astrocytes
Up in CVS-N2c
Up in SAD B19
(+) regulation of feeding behavior
cellular response to histamine
maintenance of presynaptic active zone structure
response to regulation of ventricular nicotine
cardiac muscle cell membrane repolarization
neutrophil chemotaxis
leukocyte chemotaxis
positive regulation of response to external stimulus
adaptive immune response
trachea cartilage development
inflammatory response
granulocyte chemotaxis
leukocyte migration
myeloid leukocyte migration
regulation of catecholamine secretion
granulocyte migration
cell chemotaxis
regulation of dopamine secretion
catecholamine secretion
dopamine secretion
GeneRatio
Adj. p-value
Count
O
L4
Up in CVS-N2c
Up in SAD B19
peptidyl-lysine oxidation
NO-cGMP-mediated signaling pathway
cholesterol biosynthetic process
secondary alcohol biosynthetic process
heparan sulfate metabolic process
cholesterol metabolic process
alcohol biosynthetic process
steroid metabolic process
alcohol metabolic process
steroid biosynthetic process
cell-cell adhesion via plasma-membrane
molecules cell adhesion
GeneRatio
Adj. p-value
Count
Q
L2/3
Up in CVS-N2c
Up in SAD B19
nO-cGMP-mediated signaling pathway
neurotrophic factor receptor signaling
inflammatory response
cell-cell adhesion
cell adhesion
cell-cell adhesion via plasma-membrane
(+) regulation of cell junction assembly
GeneRatio
Adj. p-value
Count
R
OL
No Significant Terms

**Supplemental Figure 5. Related to Figure 5. Differentially expressed genes and gene set enrichment analysis across cell types infected by SAD B19 or CVS-N2c. A-H.** Volcano plots showing differentially expressed genes identified between CVS-N2c- and SAD B19-infected cells of each cell type. Genes enriched in SAD B19-infected cells are shown with negative fold change, whereas genes enriched in CVS-N2c-infected cells are shown with positive fold change. Thresholds are drawn at | $\log_2$(fold change) | > 1 and adjusted p-value < 0.05. **I.** Violin plot showing enrichment scores for GO inflammatory response gene set between CVS-N2c- and SAD B19-infected cells. P-value computed using the Wilcoxon rank-sum test (median=0.2518408 for SAD B19 and 0.2495136 for CVS-N2c; *$p<0.05$). **J.** Feature plot showing enrichment scores for GO inflammatory response gene set across CVS-N2c- and SAD B19-infected cells. Highest enrichment appears in astrocytes in both datasets. **K-R.** Enriched pathways in CVS-N2c- and SAD B19-infected cells based on gene set enrichment analysis, split by cell type.

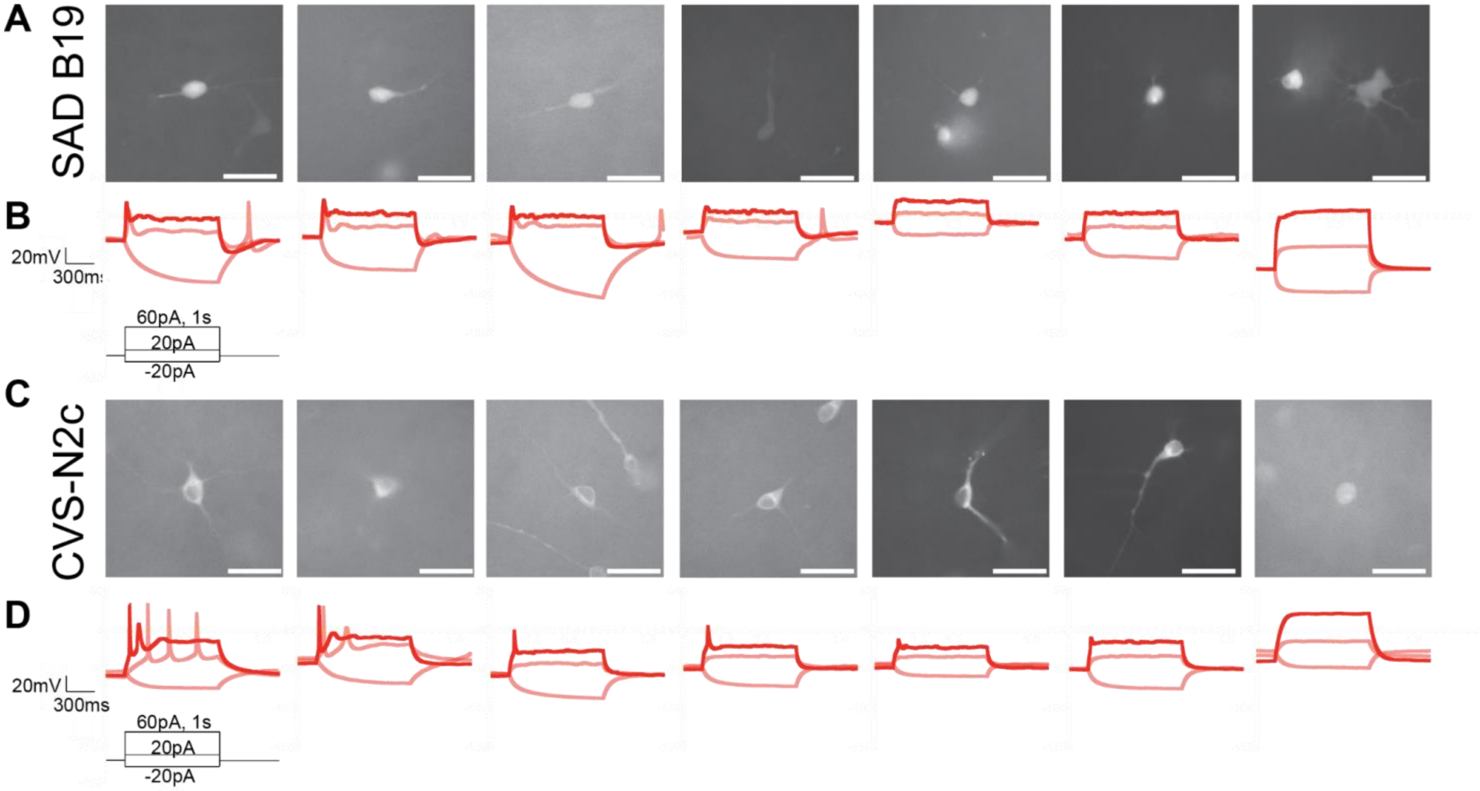


**Supplemental Figure 6. Related to Figure 5. Sample images and electrophysiological traces from patched SAD B19- and CVS-N2c-infected cells. A.** Targeted fluorescent cells infected with SAD B19 RVdG. Scale bars, 30 µm. **B.** Image-matched electrophysiological traces of SAD B19-infected neurons measured upon stimulation with the paradigm shown. **C.** Targeted fluorescent cells infected with CVS-N2c RVdG. Scale bars, 30 µm. **D.** Image-matched traces of CVS-N2c-infected neurons measured upon stimulation with the paradigm shown.

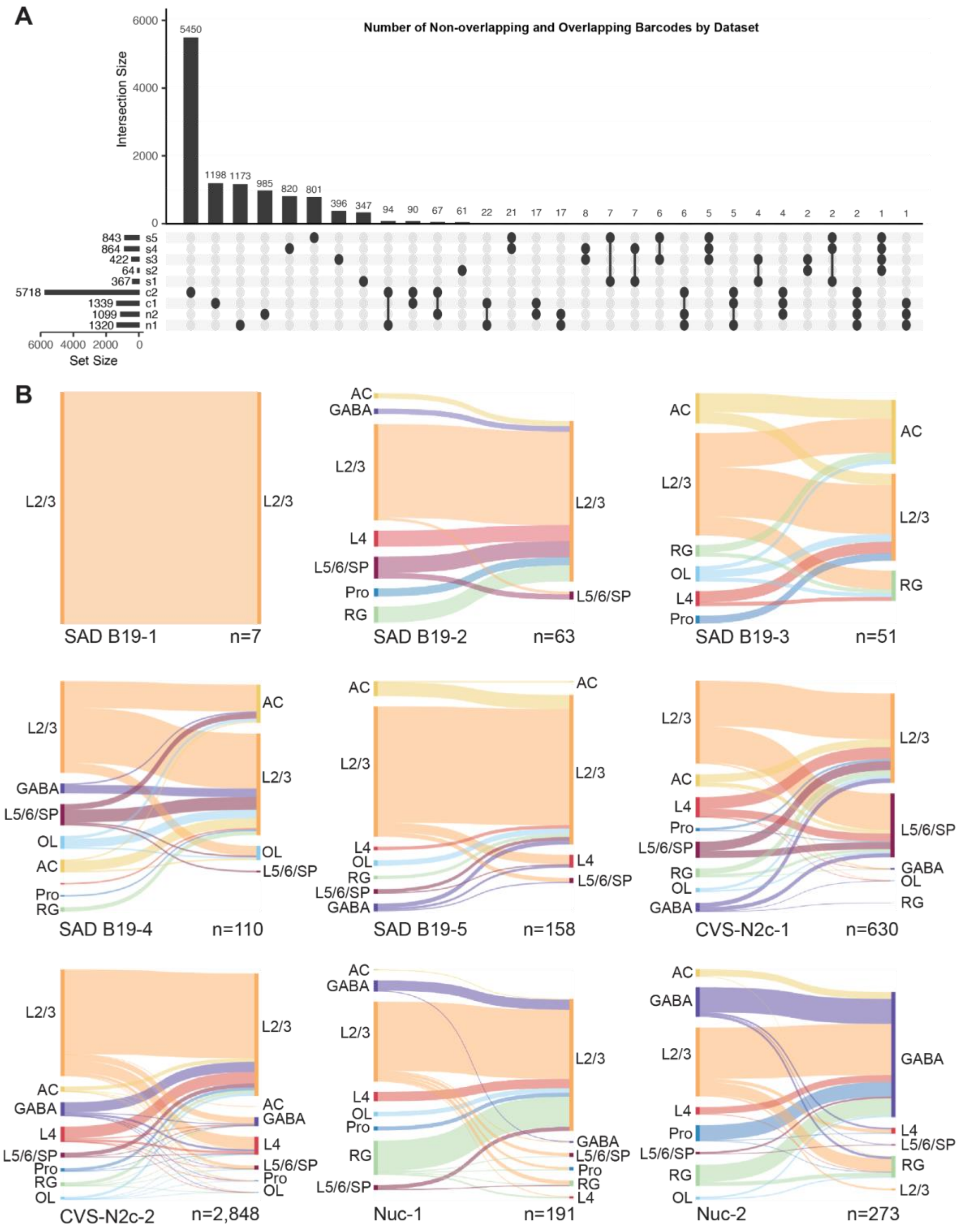


**Supplemental Figure 7. Related to Figure 8. Barcode networks across datasets. A.** Number of barcodes that are distinct or shared in each scRNA-seq or snRNA-seq dataset. **B.** Sankey plots showing directional connectivity of single starter networks across datasets. Non-starter populations are shown on the left of each plot, while starter populations are shown on the right. Number of connections observed per dataset are shown in the bottom right of each plot. Size of bars reflect the proportion of connections within that dataset.

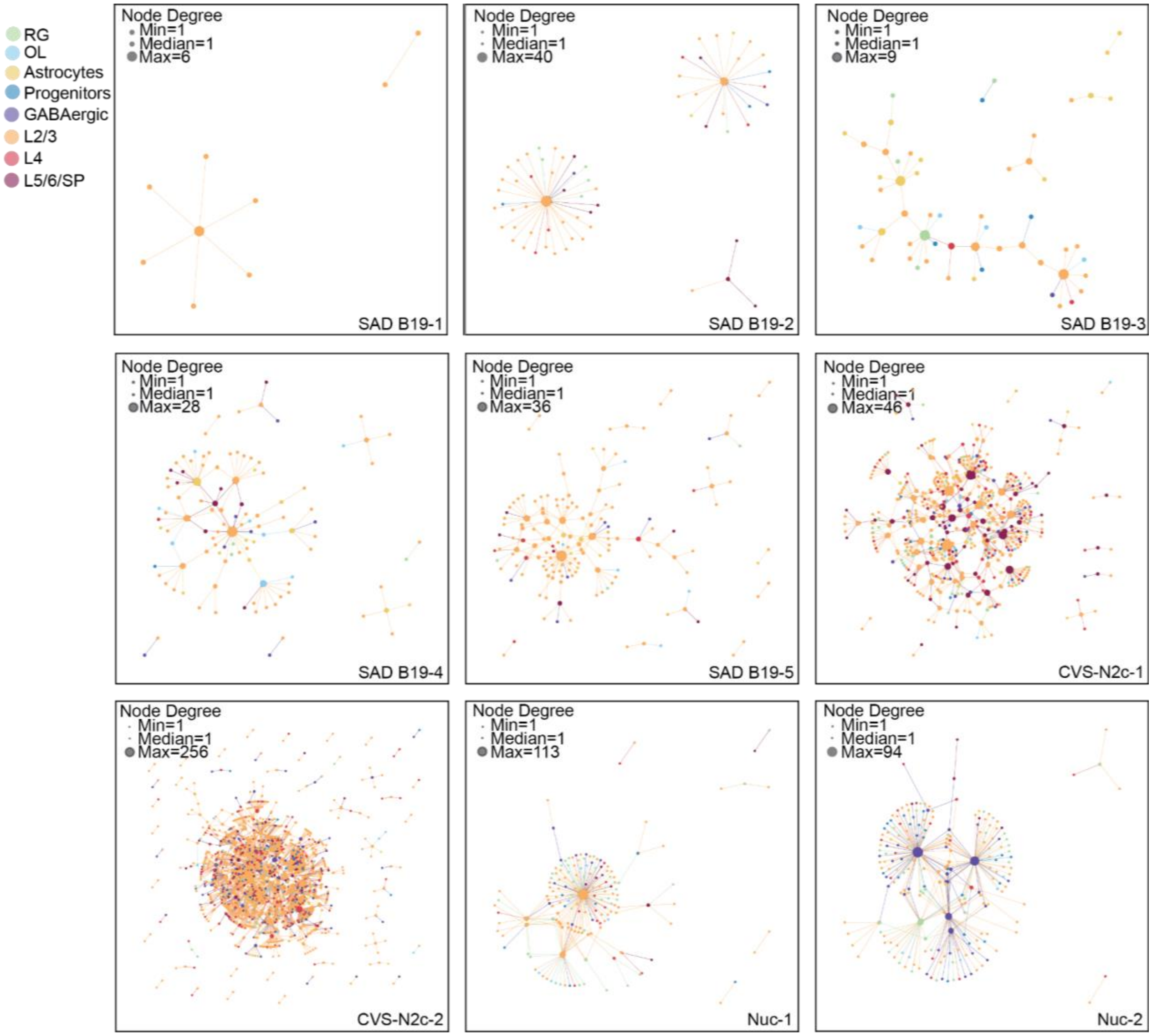


**Supplemental Figure 8. Related to Figure 8. Graph networks of connectivity across datasets.** Single starter RVdG barcode networks were plotted as graph networks after collapsing duplicate cells across barcodes. Node degrees are shown for each dataset, and cell types are annotated by color.
